# *Arabidopsis* PROTODERMAL FACTOR2 binds lysophosphatidylcholines and transcriptionally regulates phospholipid metabolism

**DOI:** 10.1101/2021.10.20.465175

**Authors:** Izabela Wojciechowska, Thiya Mukherjee, Patrick Knox-Brown, Xueyun Hu, Aashima Khosla, Graham L. Mathews, Kyle A. Thompson, Seth T. Peery, Jagoda Szlachetko, Anja Thalhammer, Dirk K. Hincha, Aleksandra Skirycz, Kathrin Schrick

## Abstract

Plant homeodomain leucine-zipper IV (HD-Zip IV) transcription factors (TFs) contain an evolutionarily conserved steroidogenic acute regulatory protein (StAR)-related lipid transfer (START) domain. The START domain is required for TF activity; however, its presumed role as a lipid sensor is not well understood. Here we used tandem affinity purification from *Arabidopsis* cell cultures to demonstrate that PROTODERMAL FACTOR2 (PDF2), a representative family member which controls epidermal differentiation, recruits lysophosphatidylcholines in a START-dependent manner. *In vitro* assays with recombinant protein verified that a missense mutation in a predicted ligand contact site reduces lysophospholipid binding. We additionally uncovered that PDF2 controls the expression of phospholipid-related target genes by binding to a palindromic octamer with consensus to a phosphate (Pi) response element. Phospholipid homeostasis and elongation growth were altered in *pdf2* mutants according to Pi availability. Cycloheximide chase experiments further revealed a role for START in maintaining protein levels, and Pi limitation resulted in enhanced protein destabilization, suggesting a mechanism by which lipid binding controls TF activity. We propose that the START domain serves as a molecular sensor for membrane phospholipid status in the epidermis. Overall our data provide insights towards understanding how the lipid metabolome integrates Pi availability with gene expression.

## INTRODUCTION

Interactions between lipids and proteins are dynamic in living organisms, yet the full extent and biological significance of such interactions is underexplored, especially in plants. In *Arabidopsis*, 21 homeodomain leucine-zipper transcription factors of the class III and IV families (HD-Zip TFs III and IV) contain a putative lipid sensor named START (Schrick et al., 2004). The steroidogenic acute regulatory protein (StAR)-related lipid-transfer (START) domain was first characterized in mammalian proteins involved in lipid transfer, metabolism and sensing (Ponting and Aravind, 1999; Alpy and Tomasetto, 2005). In humans, the START domain is found in 15 proteins, several of which are known to bind specific sterols, bile acids, phospholipids, sphingolipids, or steroid hormones (Alpy et al., 2009; Letourneau et al., 2012; Letourneau et al., 2015; Clark, 2020). Homology modeling of START domains across *Arabidopsis* HD-Zip TFs suggests that plant proteins contain a similar ligand-binding pocket (Schrick et al., 2014). In accordance, deletion of this domain from HD-Zip TF IV member GLABRA2 (GL2, AT1G79840) results in loss-of-function phenotypes that are partially complemented by the START domain from mammalian STARD1/StAR (Schrick et al., 2014). The observed complementation is abolished by a binding-site mutation, implying importance of ligand binding for GL2 function. Moreover, START domains from HD-Zip IV TFs PROTODERMAL FACTOR2 (PDF2, AT4G04890) and *Arabidopsis thaliana* MERISTEM LAYER1 (ATML1, AT4G21750), promote transcriptional activity of a chimeric TF in yeast (Schrick et al., 2014).

PDF2 and its paralog ATML1 are thought to be functionally redundant and play a critical role in maintenance of epidermal (L1) identity of the vegetative, floral and inflorescence shoot apical meristem (Abe et al., 2003). Double knockout mutants of *PDF2* and *ATML1* result in severe defects in shoot epidermal cell differentiation, resulting in embryonic lethality (Ogawa et al., 2015), while overexpression of *ATML1* is sufficient to induce epidermal identity in internal cell layers (Takada et al., 2013). Moreover, double mutants of *PDF2* with other family members (*HDG1*, *HDG2*, *HDG5*, *HDG12*), result in floral organ defects (Kamata et al., 2013a). ATML1 and PDF2 TFs bind to the L1 box, a promoter element specific to L1 genes such as those coding for extracellular proline-rich protein PROTODERMAL FACTOR1 (PDF1) (Abe et al., 2003), GDSL lipase LIP1(Rombola-Caldentey et al., 2014) and ketoacyl-CoA synthase (KCS20), the latter of which catalyzes very long chain fatty acid (VLCFA) biosynthesis (Rombola-Caldentey et al., 2014). VLCFA produced in the epidermis are thought to function as signals affecting proliferation of internal tissues *via* inhibition of cytokinin synthesis, thus modulating plant growth (Nobusawa et al., 2013). PDF2 and ATML1 are reported to interact with DELLA proteins in regulation of cell expansion (Rombola-Caldentey et al., 2014). Upon gibberellin accumulation, DELLAs are subjected to proteolysis, releasing PDF2 and ATML1 to activate expression of L1 genes (Rombola-Caldentey et al., 2014).

Considering the key role of PDF2 and ATML1 in epidermal development, we investigated additional layers of regulation, whereby TF activity is controlled by a small molecule ligand. Based on the presence of an evolutionary conserved ligand-binding domain and their role as developmental regulators, HD-Zip START TFs were suggested to constitute a link between lipid metabolism and plant development (Ponting and Aravind, 1999; Schrick et al., 2004). One prediction is that START, by binding lipids, controls gene expression analogously to steroid hormone receptors from animals. An advantage of such a mechanism is that the metabolic state of the cell would be linked to cell growth and differentiation. However, the identities of small molecule ligands of the START domain have remained elusive. To address this gap in knowledge we applied a tandem affinity purification protocol adapted for concurrent analysis of small molecule and protein partners of PDF2, a representative HD-Zip START TF from *Arabidopsis*. We then performed an *in vitro* assay that indicates direct binding of START to lysophosphatidylcholines. Our additional findings from analysis of transcriptional targets and lipidomic analysis link PDF2 TF function with phospholipid metabolism. We propose a role for PDF2 in sensing membrane phospholipid status via its START domain.

## RESULTS

### START domain of PDF2 recruits lysophospholipids *in vivo*

To investigate binding partners of the START domain from PDF2 we used tandem affinity purification (TAP) adapted for parallel analysis of protein and metabolite interactors of the bait protein of choice (Luzarowski et al., 2017; Luzarowski et al., 2018) (Figure 1A and 1B). We generated *Arabidopsis* cell lines expressing either full-length PDF2 or mutants lacking the START domain (pdf2^ΔSTART^) under control of the constitutive CaMV 35S promoter, with a TAP tag fused to either the amino- or carboxyl end. Whole cell native protein lysates (referred to as input) from cultures expressing PDF2, pdf2^ΔSTART^ or empty vector were ultracentrifuged to deplete cellular membranes. TAP-tagged proteins were immunoisolated from soluble fractions, and following stringent washes, bait proteins together with interactors were released. The eluate was extracted yielding protein pellets, polar and nonpolar (lipid) metabolite fractions.

**Figure 1.**
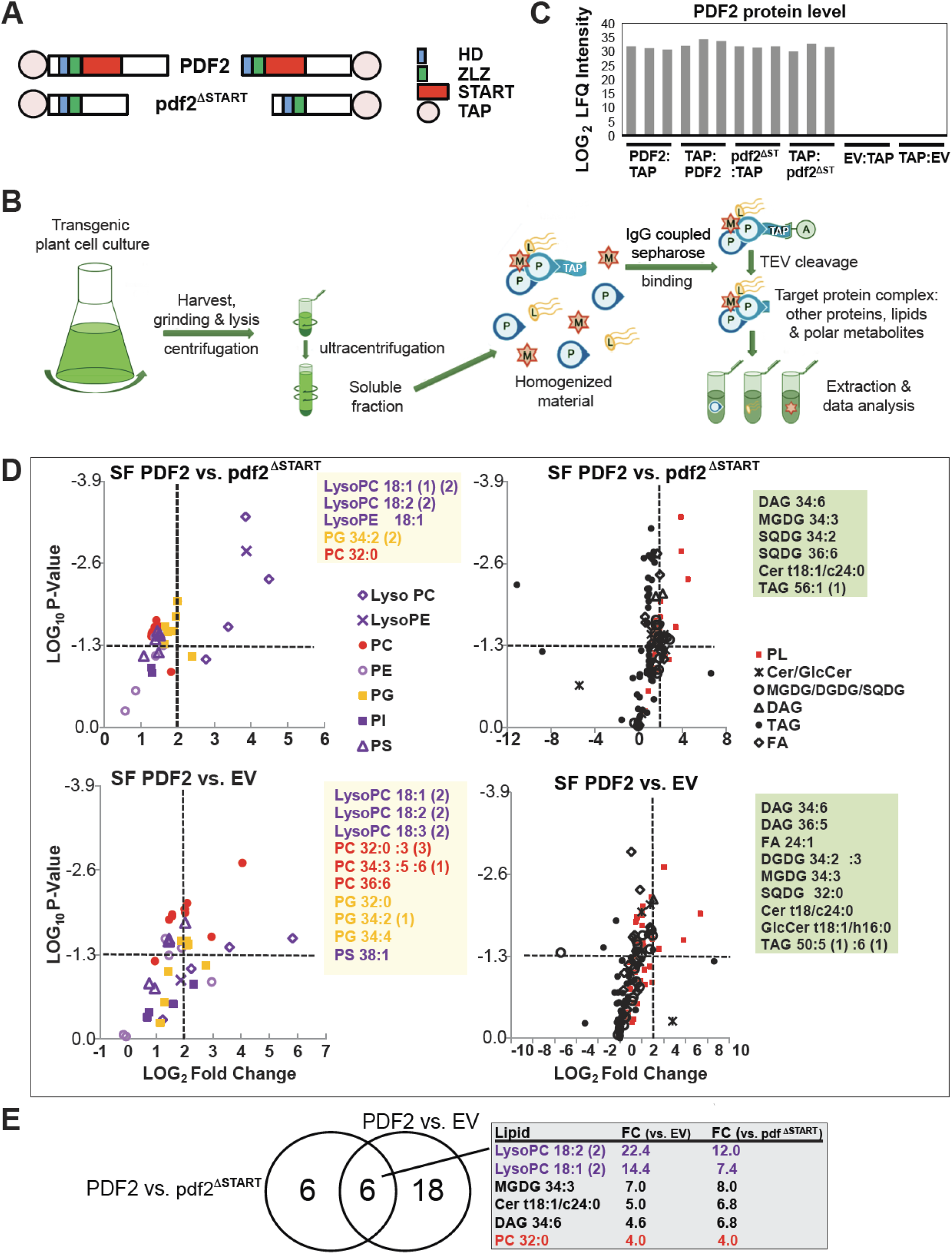
PDF2 START binds lysophosphatidylcholines in *Arabidopsis* cell cultures. **(A)** PDF2 and pdf2^ΔSTART^ proteins used for tandem affinity purification (TAP) experiments. HD, Homeodomain; ZLZ, Zipper Loop Zipper, a plant-specific leucine zipper; START domain. **(B)** Schematic of TAP protocol with *Arabidopsis* cell cultures. **(C)** PDF2 protein quantification from eluates obtained from TAP. Mass-spectrometry based proteomics revealed similar label-free quantification (LFQ) intensities for cell lines expressing full-length PDF2 and mutant pdf2^ΔSTART^. No signal was detected for empty vector (EV) lines. **(D)** START domain of PDF2 recruits lysophosphatidylcholines. Lipids were extracted from TAP eluates from PDF2, pdf2^ΔSTART^ and EV lines and analyzed by LC/MS. Volcano plots depict log2-fold changes (FC) between PDF2 and either pdf2^ΔSTART^ or EV on *x* axis versus significance (P-values, unpaired *t*-test) on *y* axis for means of 6 replicates. Horizontal dotted line indicates p = 0.05. Vertical dotted line marks a ratio of 4. SF, soluble fraction. *V*olcano plots of phospholipid (PL) changes in phosphatidylcholine (PC), phosphatidylethanolamine (PE), phosphatidylglycerol (PG), phosphatidylinositol (PI), and phosphatidylserine (PS) (left). Volcano plots of lipid profiles represent values for PL, ceramides (Cer), glucosylceramides (GlcCer), digalactosyldiacylglycerols (DGDG), monogalactosyldiacylglycerols (MGDG), sulfoquinovosyl diacylglycerols (SQDG), diacylglyerols (DAG), triacylglycerols (TAG), and fatty acids (FA) (right). Lipid interactors are shown in boxes (4-FC, *t*-test, p < 0.05). **(E)** Venn diagram illustrates lipid interactors for PDF2 but not for pdf2^ΔSTART^ or EV. See **Supplemental Data Set 1.**

Presence of the bait protein was confirmed using mass-spectrometry based proteomics (Figure 1C). To delineate a list of PDF2 lipid interactors we calculated the enrichment of the different lipids in the eluate in relation to the input. Comparison of PDF2 versus pdf2^ΔSTART^ cell lines (using input normalized data) identified 12 lipid species that were at least 4-fold more abundant (*t*-test, p < 0.05; n = 6) in PDF2 versus pdf2^ΔSTART^ lines (Figure 1D**; Supplemental Data Set 1)**. Of the 12 differential lipid species, six were also at least 4-fold more abundant (*t*-test, p < 0.05; n = 6) in PDF2 versus empty vector control lines, constituting a list of high confidence lipid binders (Figure 1D), with the highest enrichment for lysophosphatidylcholines (LysoPC 18:1 and LysoPC 18:2) (Figure 1E). No differential lipid accumulation was found between the empty vector control and pdf2^ΔSTART^ lines, implying specificity of binding **(Supplemental Data Set 1)**. The TAP experiments demonstrate that the START domain of PDF2 is associated with lipids, preferentially lysophosphatidylcholines, in cell cultures.

### Recombinant PDF2 protein binds lysophosphatidylcholines

To test whether PDF2 directly binds lysophospholipids *in vitro*, a recombinant PDF2 protein containing the ∼26 kDa START domain was produced in *E. coli* **(Supplemental Figure 1A)**. Similar to mammalian STARD1/StAR (Sluchanko et al., 2016), the PDF2 START domain is highly insoluble when expressed in *E. coli*. To enhance solubility, the maltose binding protein (MBP) was fused to its amino terminus. The MBP tag was removed by TEV protease cleavage prior to binding analysis. The pdf2(START)^L467P^ protein with a missense mutation in the C-terminal α-helix of START was used as a negative control, as L467 is a predicted ligand contact site (Roderick et al., 2002) (Figure 2A). The C-terminal α-helix is conserved in START proteins from humans as well as among HD-Zip IV TFs (Figure 2A). Analogous mutations in human StAR result in congenital lipoid adrenal hyperplasia (Bose et al., 1996; Fluck et al., 2005). Homology modeling (Roy et al., 2010; Yang and Zhang, 2015) reveals structural similarity for START from PDF2 and GL2 (Figure 2B). In GL2, the analogous L480P mutation leads to loss-of-function (Figure 2C-E).

**Figure 2.**
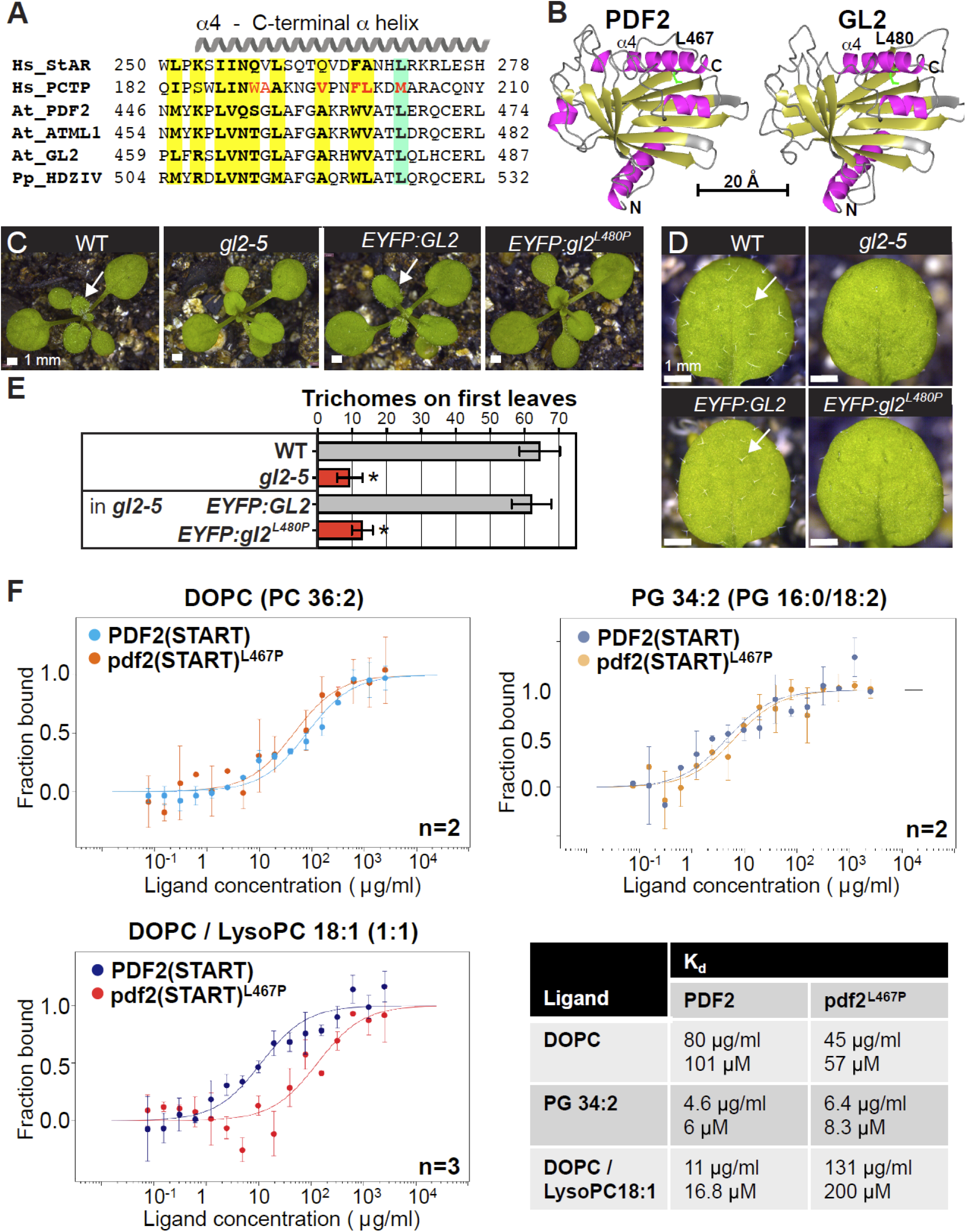
Conserved ligand contact site is required for activity and PDF2 START domain binds lysophosphatidylcholine *in vitro*. **(A)** Alignment of C-terminal α-helix of START from human (Hs) StAR and PCTP, and HD-Zip IV TFs from *Arabidopsis thaliana* (At) and *Physcomitrium patens* (Pp). Conserved amino acids (bold, yellow); conserved Leu/Met (green). Ligand contact sites as determined from PCTP-PC co-crystal (Roderick et al., 2002) (red). **(B)** Structural homology models of PDF2 and GL2 START domains generated in I-TASSER (Roy et al., 2010; Yang and Zhang, 2015) reveal conserved Leu (green) in C-terminal α-helix. **(C)** Rosettes, and **(D)** first leaves expressing *proGL2:EYFP:GL2* versus *proGL2:EYFP:gl2^L480P^* in *gl2-5* background in comparison to wild type (WT) and *gl2-5*. Normal trichomes on leaves (arrows). **(E)** Quantification of leaf trichomes: *gl2^L480P^* mutants exhibit trichome defects similar to *gl2-5*. Error bars indicate SD for *n* ≥ 20 plants. Significant differences for *gl2^L480P^* versus WT (unpaired t-test): *p < 1.0E-10. **(F)** Recombinant PDF2 START domain binds to lysophosphatidylcholines. Binding of purified PDF2(START) and pdf2(START)^L467P^ to liposomes prepared using indicated lipids, measured by microscale thermophoresis (MST). Data represented as mean ± SD of n = 2-3 independent titrations. Binding curves were used to calculate binding affinities expressed as dissociation constants K_d_. See also **Supplemental Figure 1**.

To examine binding of PDF(START) and pdf2(START)^L467P^ to lysophospholipids, we used microscale thermophoresis (MST) in conjunction with small unilamellar liposomes prepared from DOPC (36:2 PC; 1,2-dioleoyl-sn-glycero-3-phosphocholine), PG 34:2 (1-palmitoyl-2-linoleoyl-sn-glycero-3-phosphoglycerol), and a 1:1 mixture of DOPC and LysoPC 18:1 (1-oleoyl-2-hydroxy-sn-glycero-3-phosphocholine) (Figure 2F**, Supplemental Figure 1B-D)**. The data indicate that wild-type PDF(START) and mutant pdf2(START)^L467P^ bind DOPC and PG 34:2 liposomes with comparable affinities. In contrast, the presence of LysoPC 18:1 favors interaction with wild-type PDF2(START) over the mutant. Specifically, the binding affinity to DOPC/LysoPC 18:1 liposomes was ∼12-fold greater for PDF2 (START) (*K_d_* =17 µM) in comparison to pdf2(START)^L467P^ (*K_d_* =200 µM) (Figure 2F). These *in vitro* binding data indicate that PDF2 associates with and directly binds lysophosphatidylcholines through its START domain, consistent with our TAP experiments (Figure 1).

### PDF2 transcriptional targets include phospholipid catabolism genes

To investigate the connection between PDF2 and phospholipids, publicly available DNA affinity purification sequencing (DAP-seq) data (O’Malley et al., 2016) was mined for phospholipid-related gene targets using the PANTHER (Mi et al., 2019) overrepresentation test. Eight genes with gene ontology (GO) term “phospholipid catabolic process” displayed a significant enrichment of ∼9.8-fold compared to the representation expected if the target list were assembled at random (**Supplemental Data Set 2**). Putative transcriptional targets of PDF2 include non-specific phospholipase C enzymes (NPC2, AT2G26870; NPC4, AT3G03530; NPC6, AT3G48610), glycerophosphodiester phosphodiesterases (GDPD1, AT3G02040; GDPD2, AT5G41080; GDPD3, AT5G43300), and phospholipase D isoforms (PLDɛ, AT1G55180; PLDζ2, AT3G05630). Two putative targets, namely *PLD*ɛ and *PLDζ2*, were also listed as one of 29 genes with GO term “cellular response to phosphate starvation”, displaying an enrichment of ∼6.3-fold (**Supplemental Data Set 2**).

### PDF2 binds to P1BS element implicated in Pi starvation response

Genome-wide DAP-seq peak data (O’Malley et al., 2016) for PDF2 revealed the palindrome GAATATTC as the main DNA-binding motif (Figure 3A). This octamer displays consensus to the previously identified P1BS element (GNATATNC) (Rubio et al., 2001). Under Pi limitation, P1BS is the binding site of PHOSPHATE STARVATION RESPONSE 1 (PHR1), which positively regulates phospholipid remodeling and other aspects of the Pi starvation response (Pant et al., 2015).

**Figure 3.**
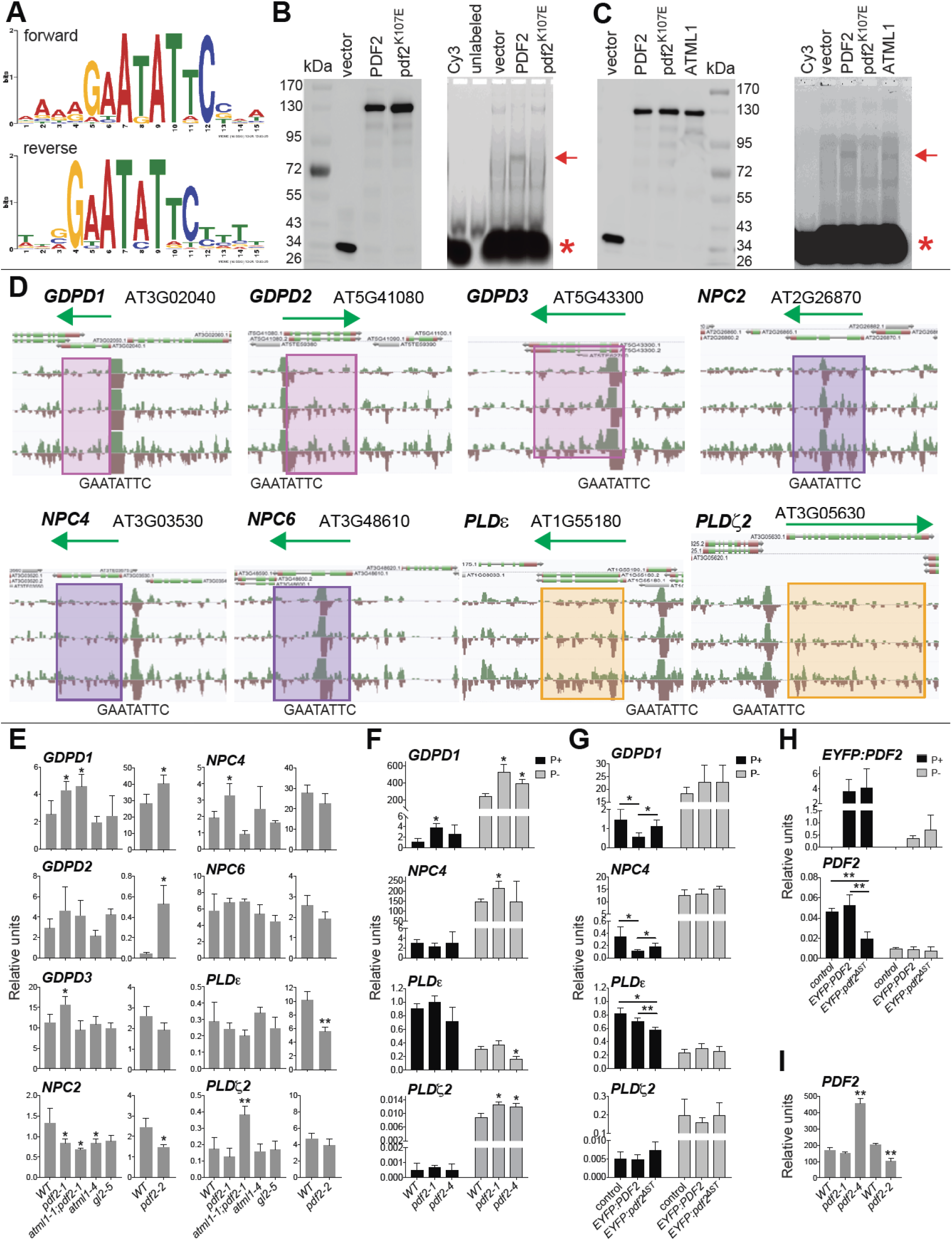
PDF2 binds a Pi response element and transcriptional targets include phospholipid catabolic genes. **(A)** Octamer motifs from DAP-seq data (O’Malley et al., 2016) for PDF2 exhibit consensus with P1BS (GNATATNC). **(B and C)** EMSA shows band shift for Halo:PDF2 (arrow), but not for HD mutant pdf2^K107E^. Asterisk indicates Cy3-labeled probe containing GAATATTC motif. Western blot with anti-Halo Ab detects Halo-tagged proteins used for EMSA (left). **(C)** EMSA shows band shift for Halo:ATML1 (arrow). **(D)** Phospholipid catabolic genes show DAP-seq peaks that map to GAATATTC on forward (green) and reverse (brown) strands. Green arrows indicate length and direction of transcript. Shaded boxes mark target gene classification: *GDPD* (pink), *NPC* (purple), *PLD* (peach). **(E)** *PDF2* is required for normal mRNA expression of several phospholipid catabolic genes. qRT-PCR with cDNA from 12 d-old (WT, *pdf2-1*, *atml1-1;pdf2-1*, *atml1-4*, *gl2-5*) or 15 d-old (WT, *pdf2-2*) seedling shoots. **(F-H)** qRT-PCR with cDNA from 14-d-old seedling shoots under Pi sufficiency (P+) or limitation (P-). **(F)** *PDF2* is required for normal gene expression of gene targets under Pi limitation. **(G)** Ectopic expression of *EYFP:PDF2* results in downregulation of *GDPD1* and *NPC4*, but upregulation of *PLD*ɛ in comparison to *EYFP:pdf2^ΔST^*. **(H)** The *EYFP:PDF2* and *EYFP:pdf2^ΔST^* lines display similar transcript levels of *EYFP* transgene. Mutant *EYFP:pdf2^ΔST^* results in downregulation of endogenous *PDF2*. **(I)** *PDF2* mRNA is upregulated in *pdf2-4* and downregulated in *pdf2-2*. **(E-I)** Data represent means of n = 4 biological replicates normalized to reference gene *ACT7*. Error bars indicate SD. Significant differences from WT or control for >3 genotypes determined by one-way ANOVA, and for 2 genotypes, unpaired *t*-test: *p < 0.05 and **p ≤ 0.0001. See **Supplemental Data Set 2**.

We used electrophoretic mobility shift assays (EMSA) to validate DNA binding of *in vitro* translated PDF2 to the P1BS palindrome. Wild-type PDF2 caused a shift in mobility of Cy3-labeled oligonucleotide containing GAATATTC (Figure 3B and 3C), in contrast to missense mutant pdf2^K107E^ in which a conserved arginine in the HD is replaced with glutamic acid (Figure 3B and 3C). Similarly to PDF2, its paralog ATML1 also bound the GAATATTC palindrome (Figure 3C). Genomic regions of eight putative target genes implicated in phospholipid catabolism were searched for the palindrome. Strikingly, P1BS elements with 100% consensus to GAATATTC overlapped with DAP-seq peaks in the promoters or 5’-UTR regions of *GDPD1, GDPD2, GDPD3*, *NPC4*, *PLD*ɛ and *PLD*ζ*2* (Figure 3D). In contrast, *NPC2* and *NPC6* exhibited peaks in internal exons of their coding regions (Figure 3D).

### PDF2 is a transcriptional repressor of several phospholipid catabolism genes

We applied quantitative real-time PCR (qRT-PCR) in conjunction with mutant analysis to test whether PDF2 is a positive or negative regulator of targets that mediate phospholipid catabolism (Figure 3E). Since PDF2 is expressed in the epidermis and the DAP-seq experiment utilized genomic DNA from young leaves (O’Malley et al., 2016), we extracted RNA from seedling shoots. This material contains epidermis as well as other tissues that do not express *PDF2*. Therefore, we considered small differences from wild type, if statistically significant, to be indicative of altered gene expression in mutants. The qRT-PCR data show that in comparison to wild-type, *GDPD1* transcripts were upregulated in *pdf2-1* and *pdf2-2*, as well as *atml1-1;pdf2-1* mutants (Figure 3E). Four of the genes (*GDPD2*, *GDPD3*, *NPC4*, *PLD*ζ*2*) also exhibited elevated transcripts in *pdf2-1*, *pdf2-2*, or *atml1-1;pdf2-1*, consistent with PDF2 acting a repressor (Figure 3E). In contrast, *atml1-4* and *gl2-5* mutants did not exhibit upregulation. The *pdf2-1*, *pdf2-2*, and *atml1-1;pdf2-1* mutants exhibited downregulation of *NPC2* (Figure 3E), while *pdf2-2* mutants showed downregulation of *PLDɛ*.

To examine the expression of selected target genes under Pi sufficiency and limitation (Figure 3F), we tested *pdf2-4* mutants carrying the null allele (Kamata et al., 2013b), alongside wild type and *pdf2-1* mutants (**Supplemental Figure 2**). Consistent with previous studies (Nakamura et al., 2005; Li et al., 2006; Cheng et al., 2011; Su et al., 2018), all genotypes showed upregulation of *GDPD1*, *NPC4*, and *PLD*ζ*2* under Pi limitation, while *PLD*ɛ was downregulated (Figure 3F). For the upregulated genes, one or both *pdf2* alleles exhibited enhanced upregulation in the case of upregulated target genes and enhanced downregulation in the case of *PLD*ɛ, a downregulated gene (Figure 3F). These results reveal that PDF2 TF activity is required for maintaining normal transcript levels of phospholipid catabolic genes under both Pi sufficiency and limitation.

We asked whether ectopic expression of *PDF2* can drive repression or activation of phospholipid catabolism target genes. We compared transgenic lines expressing *EYFP:PDF2* with mutant *EYFP:pdf2*^ΔST^ in which the START domain is deleted (Figure 3G). Wild-type *EYFP:PDF2* exhibited downregulation of *GDPD1* and *NPC4* in comparison to *EYFP:pdf2*^ΔST^ (Figure 3G). Consistent with our mutant analysis (Figure 3E and 3F), this result indicates that PDF2 is a transcriptional repressor of *GDPD1* and *NPC4*. In contrast, *PLDɛ* was upregulated in *EYFP:PDF2* in comparison to *EYFP:pdf2*^ΔST^, indicating positive regulation. The difference in activity between the wild-type and mutant transgenic lines cannot be attributed to mRNA expression since both expressed similar levels of *EYFP:PDF2/pdf2*^ΔST^ transcript (Figure 3H). Strikingly, the *EYFP:pdf2*^ΔST^ line exhibited ∼2-fold lower levels of endogenous *PDF2* mRNA (Figure 3H), consistent with the idea that this mutant, which retains the HD, interferes with autoregulation of *PDF2* through the L1 box. While *pdf2-1* showed wild-type levels of PDF2 transcript, the *pdf2-4* null allele exhibited a ∼3-fold increase in mRNA (Figure 3I). In contrast, the *pdf2-2* allele which retains the HD (Peterson et al., 2013) (**Supplemental Figure 2**) resulted in ∼2-fold lower levels of expression similar to the *EYFP:pdf2*^ΔST^ transgenic line.

### Lipidomic profiling of mutants reveals defects in phospholipid homeostasis

Gene expression changes in phospholipid catabolic genes are expected to result in altered phospholipid profiles and defects in membrane lipid remodeling. We performed a lipidomic analysis from the same shoot tissues as those used for qRT-PCR. Our LC-MS platform targeted >240 lipid species including phospholipids (LysoPC, PC, PE, PG, PI, PS), sphingolipids (ceramides (Cer) and glucosylceramides (GlcCer)), glycolipids (DGDG, MGDG, SQDG), diacyl- and triacylglycerols (DAG, TAG), and fatty acids (FA). Representative lipids from each major class were quantified in wild type and mutants for *PDF2*, *ATML1* and *GL2*. (**Supplemental Data Set 3**; **Supplemental Figures 2 and 3**). The *atml1-1;pdf2-1* double mutants display morphological defects at the seedling stage (**Supplemental Figure 2B**) (Abe et al., 2003), and we detected striking lipid changes in comparison to wild type (**Supplemental Figure 2C**), including significant differences in >100 lipid species (**Supplemental Data Set 3; Supplemental Table 1**). Phospholipids LysoPC, PE, PI, and PS were generally increased in *atml1;pdf2*. Other lipids that showed increases included DAGs, TAGs, FA, and Cer, while GlcCer, DGDG, MGDG, and SQDG were decreased (**Supplemental Data Set 3; Supplemental Figure 2C**). The other HD-Zip mutants displayed phospholipid defects to a lesser degree. The *pdf2-1* single mutants exhibited increases in several PC and PS lipids (**Supplemental Data Set 3**; **Supplemental Figure 3**). Both *atml1* alleles exhibited abnormal decreases in PG and PI lipids (**Supplemental Data Set 3**; **Supplemental Figures 2C and 3**). We also detected phospholipid alterations in *gl2-5* mutants, such decreases in PS lipid species (**Supplemental Data Set 3**; **Supplemental Figure 3**). Alterations in DAG, TAG, glycolipids and FA were additionally observed, as expected from membrane lipid remodeling.

### START domain is critical for lipid homeostasis

In a second lipidomics experiment we monitored lipid composition under Pi sufficiency and limitation for *pdf2*, *atml1* and *gl2* mutants in comparison to wild type (**Supplemental Data Set 4;** Figure 4A). To address the role of the START domain in lipid homeostasis, we included transgenic lines that were either wild-type (*PDF2* and *GL2*) or mutant for the START domain (*pdf2*^ΔST^ and *gl2^L480P^*). These transgenes are expressed as EYFP-tagged proteins in the *gl2-5* background under the epidermal-specific *GL2* promoter, which drives expression in specialized epidermal cell types including trichomes (Khosla et al., 2014). We included three *pdf2* alleles (*pdf2-1, pdf2-2* and *pdf2-4*). Based on the position of their T-DNA insertion (**Supplemental Figure 2A**), the *pdf2-1* and *pdf2-2* alleles affect START domain activity while retaining the HD, in contrast to the *pdf2-4* null allele (Kamata et al., 2013b). Likewise, *atml1-4* affects START but not HD, while *atml1-3* represents a null allele (**Supplemental Figure 2A**).

**Figure 4.**
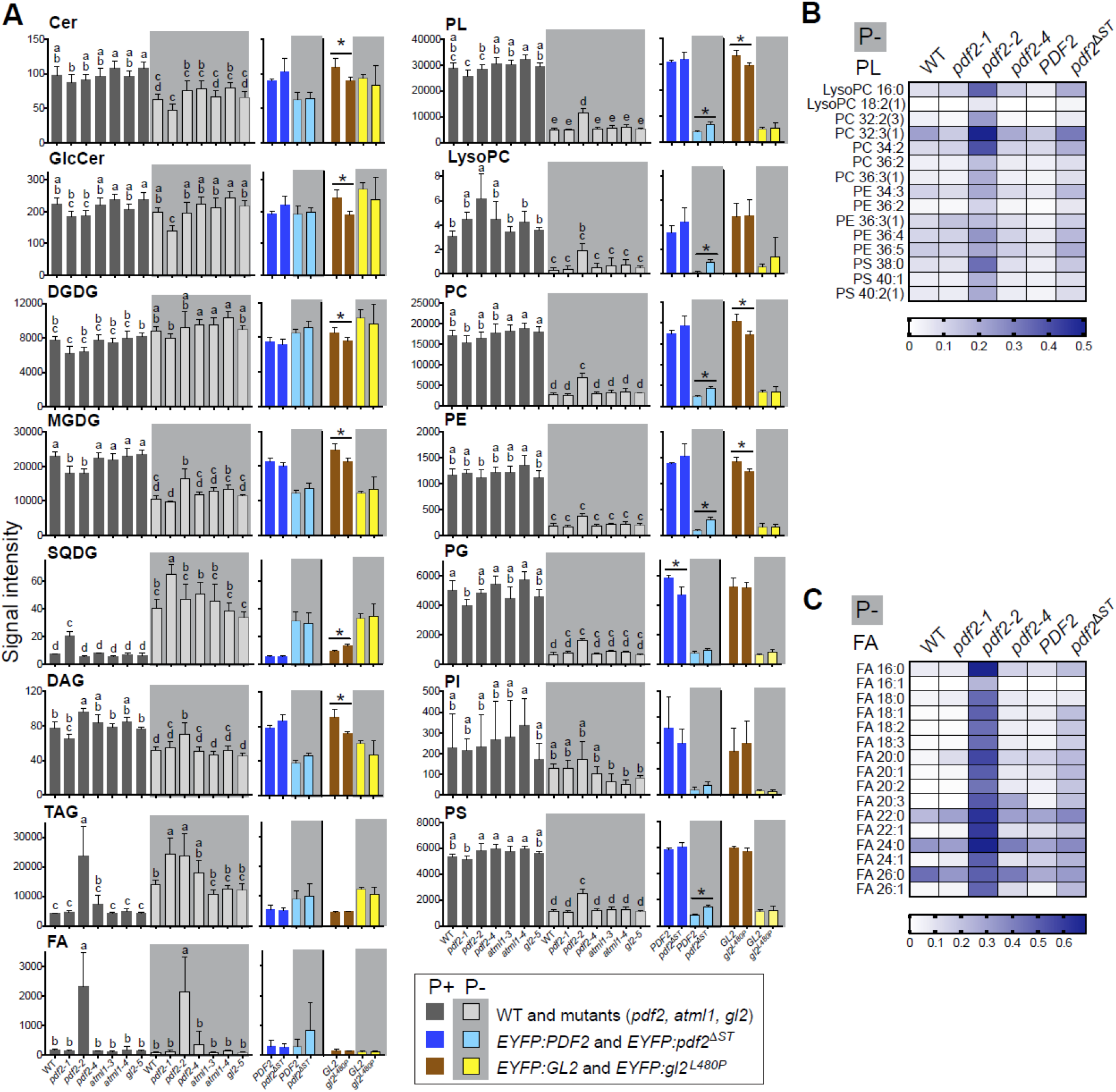
Lipidomic profiling reveals elevated phospholipid levels in *pdf2-2* and *pdf2^ΔST^* mutants under Pi limitation. **(A-C)** Lipids were extracted from 14-d-old seedling shoots from wild-type (WT), *pdf2*, *atml1*, and *gl2*, as well as *EYFP:PDF2*, *EYFP:pdf2^ΔST^*, *EYFP:GL2* and *EYFP:gl2^L480P^* transgenic lines, followed by LC-MS. Data represent 4-5 biological replicates for each genotype under Pi sufficiency (P+) or limitation (P-). Parentheses next to lipid species indicate alternative combinations of fatty acid chains corresponding to nomenclature for numbers of carbons and double bonds. **(A)** Signal intensities are indicated for means of lipid classes. Error bars indicate SD. Significant differences between >3 genotypes are marked by letters (one-way ANOVA, Tukey’s test, and between 2 genotypes by unpaired *t*-test: *p < 0.05) and **p ≤ 0.0001. **(B and C)** Heat maps of selected phospholipids (PL) **(B)** and all FA **(C)** levels under Pi limitation in WT and *pdf2* mutants in comparison to *EYFP:PDF2* and *EYFP:pdf2*^ΔST^. Only phospholipid species with significant increases in *pdf2^ΔST^* after FDR analysis are shown in (**Supplemental Table 3**) the PL heat map. Minimum and maximum values normalized to 0.0 and <1.0, respectively, for visualization purposes. See **Supplemental Figures 2-5 and Supplemental Data Sets 3 and 4**.

Pi limitation resulted in lower levels of phospholipids in wild type, as previously reported (Li et al., 2006), and we observed this trend in all lines (**Supplemental Data Set 4**; Figure 4A**; Supplemental Figure 4**). The *pdf2-2* seedlings exhibited a notably altered phospholipid profile: LysoPCs were significantly increased and others (PC, PG, PS) were increased or decreased under Pi sufficiency, whereas >30 phospholipids (LysoPC, PC, PE, PG, PS) exhibited enhanced accumulations FC ≥ 2 under Pi limitation (Figure 4A**; Supplemental Figure 5**). The *pdf2-1* mutants also exhibited altered levels of several PCs, as well as other abnormal lipid accumulations, especially TAGs and FAs, similar to *pdf2-2*, and strikingly, these defects were more pronounced under Pi limitation (Figure 4A-C; **Supplemental Table 2**; **Supplemental Table 3**).

We compared lipid profiles of seedlings expressing wild-type *EYFP:PDF2* to the *EYFP:pdf2*^ΔST^ mutant. Strikingly, *pdf2*^ΔST^ exhibited FC ≥ 2 increases in LysoPCs (16:0, 18:2, 18:3) and several other phospholipids under Pi limitation (Figure 4B**; Supplemental Figure 5; Supplemental Table 4**). In contrast, several ceramides were increased only under Pi sufficiency (**Supplemental Figure 5**). Similarly, when we compared wild-type *EYFP:GL2* to START domain mutant *EYFP:gl2^L480P^* we also noted lipid changes that varied with Pi status (**Supplemental Figure 5H**). LysoPC 18:2 and several FAs were elevated FC ≥ 2 in *gl2^L480P^* under Pi limitation. We compared lipid changes in *pdf2-2* and *pdf2*^ΔST^ which both affect START domain (but not HD) function and both exhibit reduced levels of endogenous *PDF* transcript (Figure 3H and 3I). Under Pi limitation, *pdf2-2* and *pdf2*^ΔST^ shared numerous phospholipid increases in comparison to controls (Figure 4B**; Supplemental Figure 5; Supplemental Table 3; Supplemental Table 4**). We also noted a trend indicating multiple FA species elevated in both START domain mutants (Figure 4C). Overall, the results suggest imbalances in phospholipid and FA levels in START domain mutants, notably under Pi starvation.

### *PDF2* drives elongation growth in the root

To identify a growth phenotype associated with HD-Zip function, we assayed null mutant seedlings for *PDF2*, *ATML1* and *GL2* for vertical root growth in Pi sufficient and limiting media (Figure 5A). The *pdf2-4* null mutant exhibited altered elongation growth under Pi sufficient conditions, while *atml1-3* and *gl2-5* null mutants appeared indistinguishable from wild type (Figure 5B). Root elongation was mildly altered for all three null mutants (*pdf2-4*, *atml1-3* and *gl2-5*) under Pi limitation (Figure 5B).

**Figure 5.**
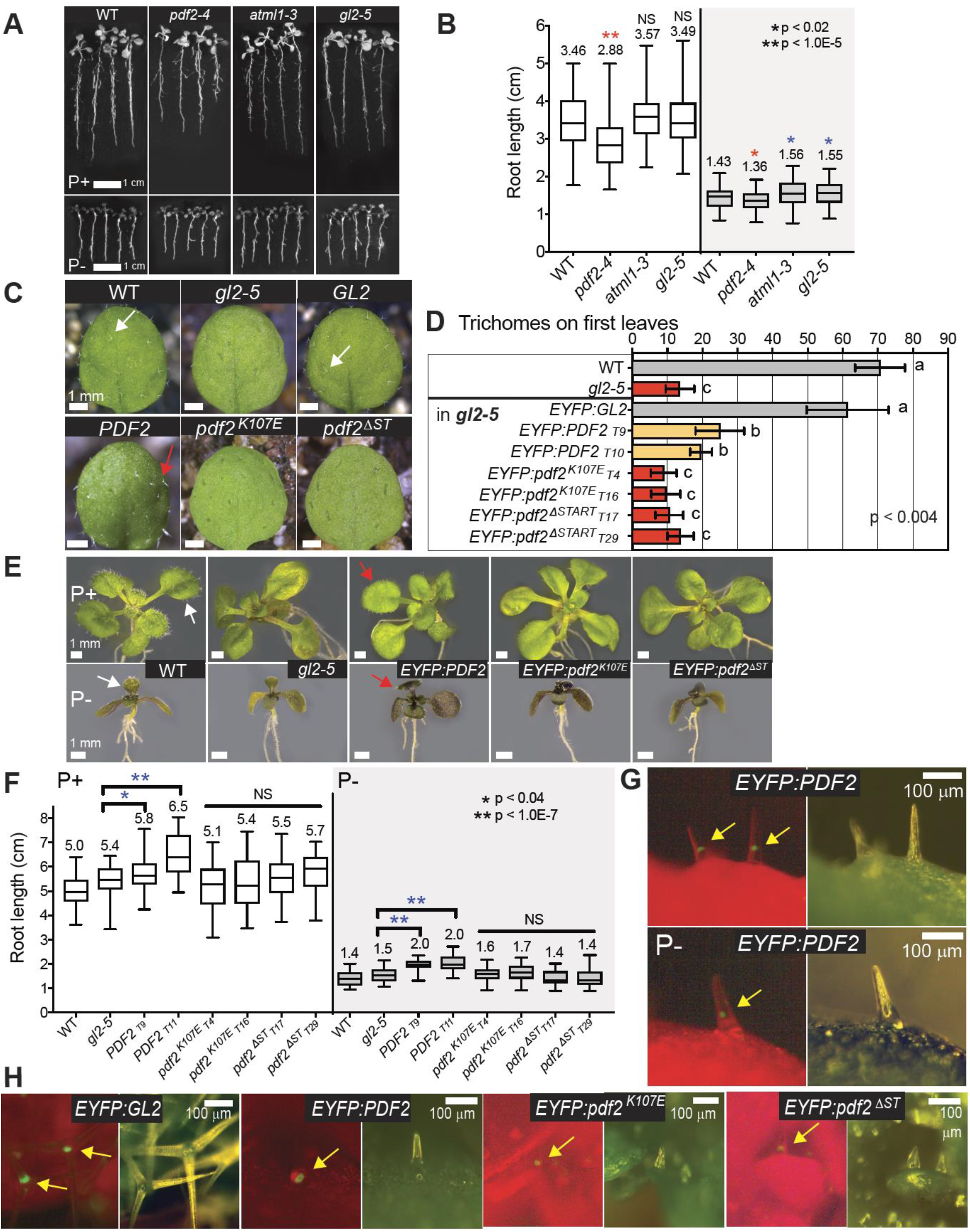
START domain-dependent expression of PDF2 drives elongation growth. **(A)** Seedlings from wild type (WT), and HD-Zip IV null mutants were grown under Pi sufficiency (P+) or limitation (P-) for 14 d. Size bars = 1.0 cm. **(B)** Root lengths for seedlings from each genotype. Each box plot represents n > 130 seedlings from 4-5 independent experiments. Horizontal lines denote median. Vertical lines indicate minimum and maximum values. Means are reported above box plots. Significant decreases (red) or increases (blue) from WT (unpaired *t*-test): *p < 0.02 or **p < 0.00001. **(C)** Ectopic expression of PDF2 results in gain-of-function trichome phenotype that is HD and START domain dependent. First leaves of WT and *gl2-5* plants in comparison to *proGL2:EYFP:GL2*, *proGL2:EYFP:PDF2*, *proGL2:EYFP:pdf2^K107E^* and *proGL2:EYFP:pdf2^ΔST^* in *gl2-5* background. Arrows indicate normal (white) and abnormal (red) leaf trichomes. Size bars = 1 mm. See also **Supplemental Figure 6.** **(D)** Quantification of leaf trichomes. Error bars indicate SD for n ≥ 20 plants (unpaired t-test): *p < 0.00001). **(E)** Phenotypes of 14-d-old seedlings under P+ or P-conditions. Arrows indicate normal (white) and abnormal (red) leaf trichomes. Size bars = 1 mm. **(F)** Ectopic expression of PDF2 drives root elongation. Quantification of root lengths for seedlings shown in **(E)**. Each box plot represents n ≥ 29 seedlings from two independent experiments. Two independent transformants (T#) were analyzed for each transgene. Significant increases (blue) to control (unpaired *t*-test): *p < 0.04 or **p < 1.0E-7. **(G and H)** Epifluorescence (left) with matching light images (right) of leaf trichomes from 14-d-old seedlings. Size bars = 100 µm. **(G)** EYFP:PDF2 is nuclear localized under P+ and P-conditions (arrows). **(H)** EYFP-tagged wild-type and mutant proteins are nuclear localized (arrows).

We next examined seedlings expressing *EYFP:PDF2* under the epidermal specific *GL2* promoter in the *gl2-5* background. The *GL2* promoter drives expression in trichomes and in non-root hair cells (Khosla et al., 2014), which undergo extensive elongation in the seedling. Expression of wild-type *EYFP:PDF2*, but not HD mutant *EYFP:pdf2^K107E^* or START mutant *EYFP:pdf2*^ΔST^, partially rescued the trichome defect of *gl2-5* (Figure 5C and 5D). To further test whether *PDF2* is critical for elongation growth we measured root lengths in *EYFP:PDF2* versus *EYFP:pdf2*^ΔST^ seedlings (Figure 5E and 5F). The data indicate that ectopic expression of wild-type *PDF2* under both Pi sufficiency and limitation results in increased elongation, whereas elongation in *pdf2^K107E^* or *pdf2*^ΔST^ was indistinguishable from the control. At later stages, we observed growth defects and aberrant leaf morphologies in the *PDF2* expressing lines, but not in *pdf2^K107E^* or *pdf2*^ΔST^ lines (**Supplemental Figure 6**). EYFP-tagged PDF2 protein exhibited nuclear localization under both Pi sufficiency and limitation (Figure 5G), and mutant pdf2 proteins were also expressed in nuclei (Figure 5H). The data indicate that ectopic epidermal *PDF2* expression drives elongation growth in the root, and the observed growth phenotype is dependent on both the HD and START domains.

### START domain mutation L480P affects elongation growth and repression of ***PLDζ1***

We further tested whether the START domain is required to control elongation growth by comparing *gl2-5* seedlings stably expressing *proGL2:EYFP:GL2* or *proGL2:EYFP:gl2^L480P^*. The START domain mutation L480P leads to trichome defects (Figure 2C-2E; Figure 6A). Additionally, the *gl2^L480P^* seedlings displayed slightly decreased elongation under Pi sufficiency, and increased elongation in comparison to wild type under Pi limitation (Figure 6B). Although both transgenes were expressed and the respective proteins were nuclear localized, only wild-type *GL2* but not mutant *gl2^L480P^* showed repression of phospholipase target gene (Ohashi et al., 2003) *PLD*ζ*1* (Figure 6C and 6D). The differential growth phenotype of *gl2^L480P^* versus wild-type *GL2* as well as qRT-PCR analysis suggests that a functional ligand-binding START domain is critical for normal elongation growth and target gene repression in response to Pi availability.

**Figure 6.**
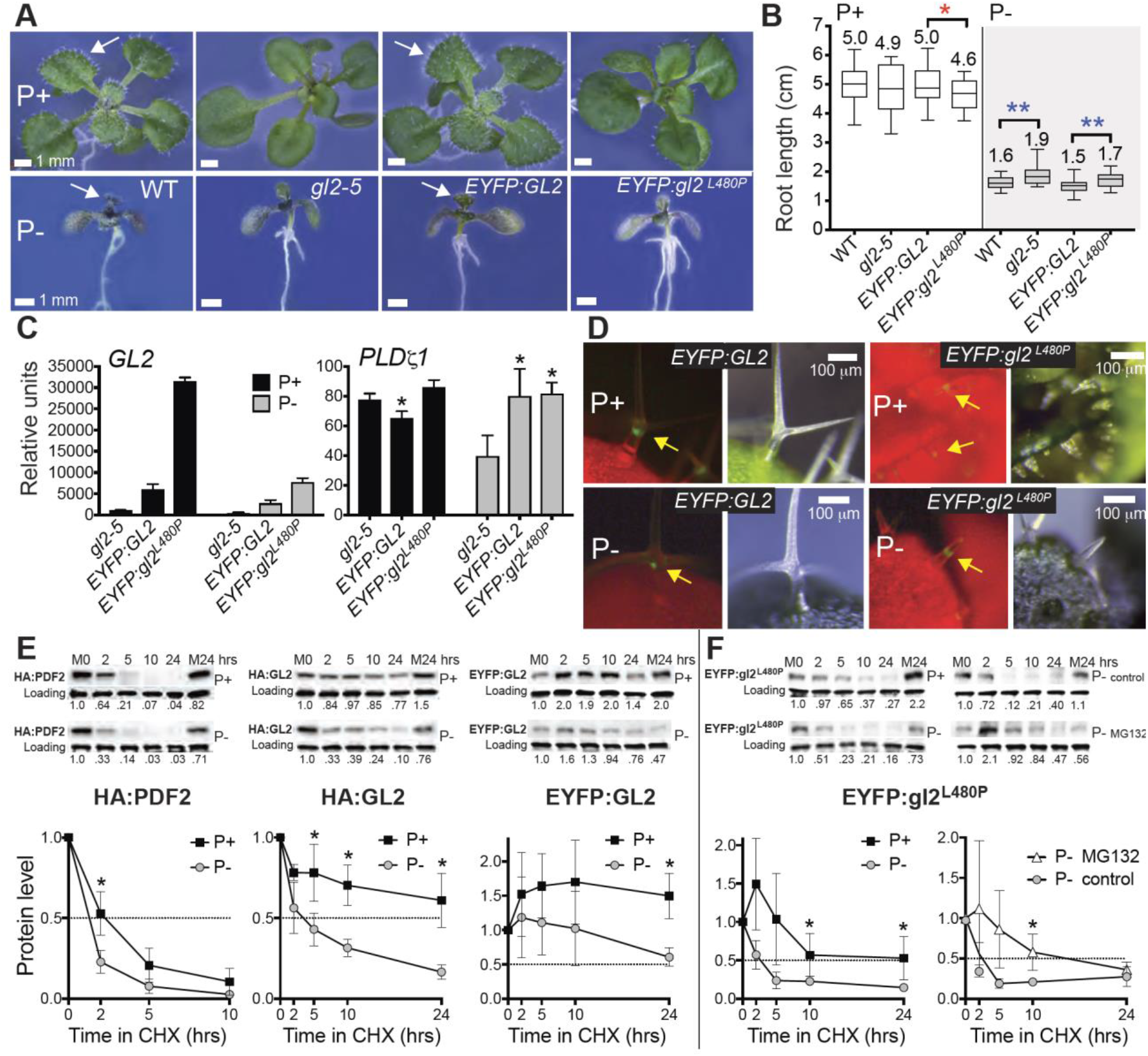
START mutant L480P affects root elongation, target gene repression, and protein stability of GL2. **(A)** Phenotypes of 14-d-old seedlings from wild type (WT), *gl2-5*, *proGL2:EYFP:GL2* and *proGL2:EYFP:gl2^L480P^* (in *gl2-5* background) under Pi sufficiency (P+) or limitation (P-). Normal leaf trichomes (arrows). Size bars = 1 mm. **(B)** Root lengths for n ≥ 21 seedlings. Significant decrease (red) or increase (blue) to control (unpaired *t*-test): *p < 0.04 or **p < 0.002. **(C)** qRT-PCR with cDNA from 14-d-old seedling shoots under P+ or P-conditions, normalized to reference gene *ACT7*. Both WT and mutant lines express *GL2/gl2* transcript. *EYFP:GL2* but not *EYFP:gl2^L480P^* exhibits repression of *PLDζ1* under P+ conditions. Significant difference to *gl2-5* (unpaired *t*-test): *p < 0.05. **(D)** Epifluorescence (left) with matching light images (right) of leaf trichomes from 14-d-old seedlings. EYFP:GL2 and EYFP:gl2^L480P^ are nuclear localized (arrows) under P+ and P-conditions. Size bars = 100 µm. **(E)** Protein stability of PDF2 and GL2 is reduced under Pi limitation. Seedlings were grown on P+ or P-media for 5-6 days, followed by cycloheximide (400 µM) treatment for 24 h. Western blot with anti-HA or -GFP antibodies, followed by Coomassie blue staining for loading controls. Top (P+) and bottom (P-) rows are from the same blot. M0 and M24, DMSO mock treatments at 0 and 24 h. Each blot is representative of n = 3-4 independent experiments. **(F)** In comparison to GL2 **(E)**, gl2^L480P^ exhibited a shorter half-life that was enhanced under Pi limitation. MG132 (50 µM) treatment restored gl2^L480P^ stability. **(E and F)** Protein quantification, with values normalized to M0, are graphed beneath Western blots. Intersection with dotted line (0.5) denotes protein half-life. Error bars indicate SD. Significant differences (unpaired t-test): *p ≤ 0.05. See also **Supplemental Figure 7.**

### PDF2 and GL2 exhibit reduced protein stability under Pi limitation, and protein destabilization is enhanced in START mutant L480P

Time-course microarray profiles of *Arabidopsis* seedlings previously indicated that *PDF2*, *ATML1* and *GL2* transcripts are not significantly up- or downregulated in the initial response to Pi starvation (Lin et al., 2011). However, our qRT-PCR data indicate that prolonged Pi limitation results in downregulation of both *PDF2* and *GL2* transcripts in seedlings (Figure 3), possibly due to feedback mechanisms affecting TF function. Therefore, we asked whether HD-Zip TF levels are post-translationally regulated. We performed cycloheximide assays with seedlings expressing tagged TFs to determine whether the START domain affects protein stability depending on Pi status. We first examined the stability of hemagglutinin-tagged proteins, HA:PDF2 and HA:GL2, and found both to exhibit reduced half-lives under Pi limitation (Figure 6E**; Supplemental Figure 7**). We next examined the EYFP:GL2 protein and found it to be stable over a 24-h time course under Pi sufficiency (Figure 6E). The increased stability of EYFP:GL2 in comparison to HA:GL2 is likely due to the larger tag (∼28 kDa versus ∼1 kDa). Similar to the HA-tagged proteins, EYFP:GL2 exhibited reduced stability under Pi limitation. In comparison, the EYFP:gl2^L480P^ mutant protein exhibited a decrease in stability and half-life of ∼10 h under Pi sufficiency (Figure 6F**; Supplemental Figure 7**). The half-life of EYFP:gl2^L480P^ was further reduced to ∼2 h under Pi limitation (Figure 6F), indicating that START is critical for protein stability under both conditions. Coincubation of seedlings with cycloheximide and proteasome inhibitor MG132 restored stability of EYFP:gl2^L480P^ under Pi limitation (Figure 6F), suggesting that EYFP:gl2^L480P^ protein is degraded via the 26S proteasome. These experiments reveal that HD-Zip TFs are destabilized under Pi limitation, and that the START domain contributes to protein stability.

## DISCUSSION

### HD-Zip protein PDF2 binds lysophospholipids via its START domain

The main finding herein is that PDF2, via its START domain, directly interacts with lysophosphatidylcholines. Our initial strategy was to identify *in vivo* binding partners of this representative HD-Zip TF by performing TAP experiments with *Arabidopsis* cell lines. We followed up on candidate ligands using *in vitro* binding validation. Our data are consistent with a previous study in which START domains of PDF2, ATML1, and GL2 were heterologously expressed in yeast and subjected to immunoisolation (Schrick et al., 2014). Subsequent lipidomic analysis revealed enrichment of lysophosphatidylcholines and other phospholipids (PC and PS) in START domain pull-down samples (Schrick et al., 2014). Although it is possible that the epidermal cells in which these HD-Zip TFs are predominantly expressed contain additional ligands, lysophospholipids now emerge as important PDF2 interactors.

Lysophosphatidylcholine arises from partial hydrolysis of PC to remove one of the fatty acid groups. Since Pi starvation induces breakdown of PC in plants, lysophosphatidylcholines serve as intermediates of the plastidic lipid biogenesis pathway. It was proposed ∼20 years ago that lysophosphatidylcholine is exported from ER to chloroplast as a precursor for galactolipid synthesis (Mongrand et al., 2000). Lysophospholipids are additionally thought to serve as messengers in plants. In arbuscular mycorrhizal symbiosis, roots use lysophosphatidylcholine as a signaling molecule to induce expression of endogenous Pi transporter genes (Drissner et al., 2007).

PDF2 START domain binding to lysophosphatidylcholines in *Arabidopsis* cells (Figure 1) and in yeast (Schrick et al., 2014) as well as *in vitro* (Figure 2F) builds on mounting evidence that links HD-Zip IV TFs with phospholipid sensing. In 2003, GL2 was identified as a negative regulator of phospholipase D (*PLD*ζ*1*) in root hair patterning (Ohashi et al., 2003). Further insights came from studies with mammalian STARD2/phosphatidylcholine transfer protein (PCTP), which binds phosphatidylcholine and is expressed during embryonic development in the mouse. STARD2/PCTP interacts with and enhances TF activity of Pax3, a mammalian HD protein (Kanno et al., 2007). The START domain from human PCTP, similarly to the PDF2 START domain, also recruits lysophosphatidylcholines in pull-down experiments in yeast (Schrick et al., 2014). Our findings introduce the intriguing possibility that START-dependent mechanisms linking Pi sensing and transcriptional control of phospholipid metabolism are conserved across organisms.

### Dual role of PDF2 as a metabolic sensor and transcriptional regulator of phospholipid metabolism

Here we identify PDF2 as a negative regulator of several phospholipid catabolism genes. Until now, PDF2 was viewed as an activator that functions redundantly with ATML1 to positively regulate L1 genes. Surprisingly, the DAP-seq data identified the P1BS element (GAATATTC) as the main DNA-binding motif for PDF2 (Figure 3A), as opposed to the L1 box (TAAATCTA), which was reported as the DNA-binding motif for both ATML1 and PDF2 (Rombola-Caldentey et al., 2014). Our gene expression studies show that *pdf2*, and not *atml1* mutants exhibit transcriptional upregulation of several phospholipid catabolism genes, suggesting that PDF2 is the main repressor of these genes in the shoot. Moreover, ectopic *PDF2* expression was sufficient to drive repression (Figure 3G). Since ATML1 also binds the P1BS element (Figure 3C), its activity may be critical for a different subset of target genes. The lipidomics data (**Supplemental Data Sets 3-5**; Figure 4; **Supplemental Figures 2-5)** implicate both ATML1 and PDF2 as regulators of phospholipid homeostasis.

We propose that PDF2 functions as a lipid sensor for phospholipids via its START domain (Figure 7A). In our model, lysophosphatidylcholines bind to START to stabilize the protein, resulting in transcriptional activity. PDF2 directly binds to the promoters of several phospholipid catabolism genes to promote incorporation of phospholipids into membranes, driving elongation growth. Mutant analysis indicates that PDF2 is important for transcript levels of phospholipid catabolic genes (Figure 3). *PLD*ɛ overexpression enhances root growth under Pi deprivation (Hong et al., 2009), consistent with our finding that PDF2 positively regulates this target gene. Under Pi starvation, overall phospholipid levels including lysophospholipids decrease, resulting in reduced PDF2 protein levels and reduced cell elongation. However, PDF2 levels are not completely abolished. According to this model, the transcriptional activity of PDF2 is critical to regulate phospholipid catabolic genes to allow measured growth according to lysophospholipid levels. Under Pi limitation, we suggest that a pool of PDF2 protein is either unliganded or bound to a destabilizing ligand, resulting in proteasome-mediated degradation.

**Figure 7.**
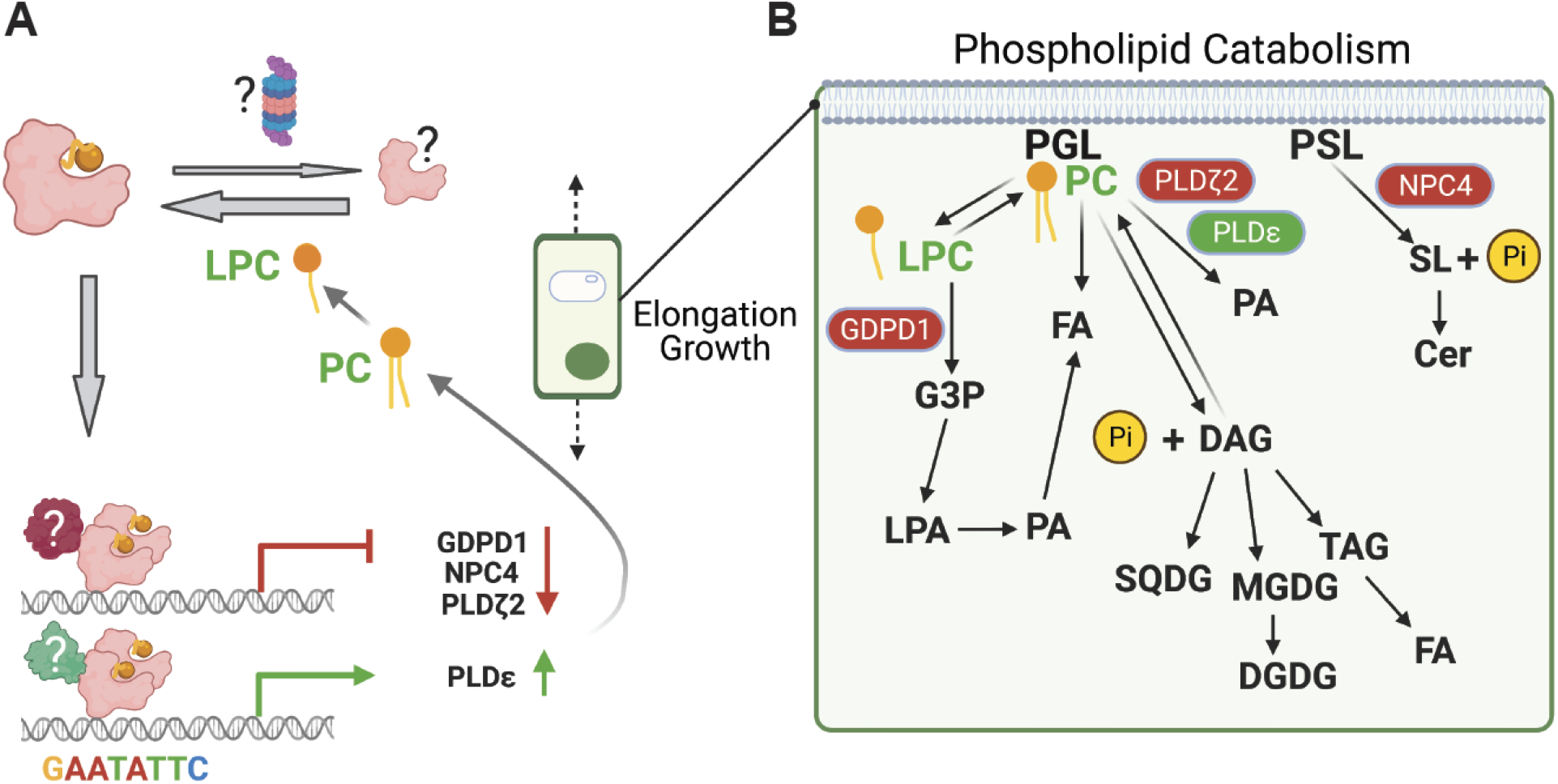
Model for the role of PDF2 as a lipid sensor. **(A)** The START domain of PDF2 binds a lipid ligand, resulting in either stabilization or destabilization of the protein, possibly via the 26S proteasome. In the illustrated scenario, lysophosphatidylcholine binding results in stabilized TF that dimerizes and binds to the P1BS palindrome upstream of phospholipid catabolic genes. Negative or positive regulation of gene expression occurs through interaction with an unknown corepressor or coactivator, respectively. This gene regulation drives the maintenance of membrane phospholipids in epidermal cells undergoing elongation growth, even under Pi limitation. **(B)** Lysophosphatidylcholine plays a central role in phospholipid catabolism. Phosphoglycerolipids (PGL) and phosphosphingolipids (PSL) of the plasma membrane are major stores of Pi in the cell. The GDPD, NPC4, and PLDζ2 enzymatic steps are transcriptionally repressed by PDF2. In contrast, PLDɛ enzyme activity, which is associated with enhanced root growth and biomass accumulation (Hong et al., 2009), is promoted by PDF2 transcriptional activation. These events result in phospholipid accumulation and production of lysophosphatidylcholine, which in turn binds PDF2 to positively regulate its activity **(A)**. This figure was created with BioRender.com.

PDF2 activity is positioned to protect membrane lipid biogenesis in the epidermis when Pi is limiting. Derepression of phospholipid catabolic genes leads to the production of fatty acids, glycolipids, as well as DAG and TAG, and recycling back to phospholipids (Figure 7B). Our lipidomic profiling of *pdf2*, *atml1*, and *gl2* mutants uncovered altered levels of several types of phospholipids, as well as products of phospholipid catabolism. While *pdf2* null mutants exhibited elongation defects in the seedling, we found that ectopic expression of *PDF2* drives root elongation. The growth promoting activity in the seedling requires the function of both the START domain and HD (Figure 5F). In contrast, ectopic expression of *EYFP:PDF2* or *EYFP:ATML1* under the *GL2* epidermis-specific promoter leads to dwarfism in adult plants (**Supplemental Figure 6**), a phenotype that is abolished by HD or START domain mutation. These observations highlight the importance of PDF2 and ATML1 function in maintaining the normal growth pattern.

Why should a phospholipid sensing mechanism that transcriptionally controls phospholipid catabolism function in the epidermis? In addition to its myriad protective functions, the epidermis plays a critical role in controlling growth. The brassinosteroid pathway for cell expansion and cell division is required in the L1 layer (Savaldi-Goldstein et al., 2007), and epidermis-localized VLCFA biosynthesis is implicated in growth control (Nobusawa et al., 2013). *NPC4*, which is negatively regulated by PDF2 (Figure 3), was reported to be critical for hydrolysis and breakdown of glycosyl inositol phosphoceramides (GIPC) (Yang, 2021). These phosphosphingolipids, along with phosphoglycerolipids, are major constituents of the plasma membrane. We uncovered evidence that *PDF2* negatively regulates *PLDζ2*, a gene responsible for Pi-deficit induced PC hydrolysis (Su et al., 2018). It is estimated that in plants, about one-third of cellular Pi is stored in membrane phospholipids. Our study highlights the importance of membrane phospholipids and lipid homeostasis as a regulator of growth in the epidermis.

### Perspectives on START domains as phospholipid sensors

Whether other START domain-containing HD-Zip TFs besides PDF2 bind lysophospholipids needs to be tested experimentally. Considering that mammalian START proteins differ in their specificity towards various lipids ranging from fatty acids to sterols, a similar diversification is expected in plants. VLCFA-ceramides were recently proposed to bind the START domain of ATML1 (Nagata et al., 2021). Fitting with this possibility, our TAP results for PDF2 identified one ceramide species (Cer t18:1/c24:0) that is enriched in wild-type versus the *pdf2*^ΔSTART^ mutant (Figure 1E). Aside from HD-Zip III and IV TFs, *Arabidopsis* contains 14 START proteins whose ligands are unknown (Schrick et al., 2004). Recently, a START protein from *Marchantia* was implicated in lipid transfer activity during Pi deprivation (Hirashima et al., 2021). The only other plant START protein reported to bind lipids, the wheat stripe rust resistance protein WKS1, appears to show specificity towards phosphatidic acid and phosphatidylinositol phosphates in lipid blots (Gou et al., 2015).

It is noteworthy that this newly discovered lipid metabolism connection relates to sensing of Pi, a nutrient that is crucial for plant growth. Our findings open a new area of research that will further explore how Pi sensing and membrane lipid metabolism are integrated with the developmental program in plants and across multicellular organisms. Intriguingly, a human START protein of the thioesterase family (THEM1/STARD14) that is critical for brown fat metabolism is allosterically regulated via its binding to lysophosphatidylcholine in addition to fatty acids (Tillman et al., 2020). Since both lysophosphatidylcholines and fatty acids are breakdown products of membrane lipid catabolism in plants, future avenues of research will explore how START domains evolved to effectively orchestrate gene expression networks according to environmentally guided metabolic inputs.

## METHODS

### Plant cell cultures, Plants and growth conditions

PSB *Arabidopsis thaliana* cell cultures (Van Leene et al., 2011) were grown in MSMO medium with 3% sucrose, 0.05 mg/L kinetin and 0.5 mg/L 1-naphthaleneacetic acid at 130 rpm. Cells were passaged weekly to fresh medium and harvested during logarithmic growth using rapid filtration and liquid nitrogen snap freezing. Transformation with TAP constructs was as described previously (Van Leene et al., 2011). *Arabidopsis thaliana* plants were of the Columbia (Col-0) ecotype. Seeds for *pdf2-1* and *atml1-1* (Abe et al., 2003) were provided by Taku Takahashi. Both *atml1-3* (SALK_033408) and *atml1-4* (SALK_128172) are T-DNA insertion alleles (Roeder et al., 2012) provided by Adrienne Roeder. *gl2-5* is a En-1 insertion allele of *GL2* (Ohashi et al., 2003; Khosla et al., 2014). *pdf2-2* (SALK_109425) and *pdf2-4* (SAIL_70G06) T-DNA insertion lines (Kamata et al., 2013b; Peterson et al., 2013) were from ABRC. Genotyping primers are listed in **Supplemental Table 5**. *HA:PDF2* and *HA:GL2* were transformed into Col-0 plants. The *proGL2:EYFP:GL2, proGL2:EYFP:PDF2* and *proGL2:EYFP:PDF2* constructs (and mutant variants) were transformed into *gl2-5*, while *proGL2:EYFP:PDF2* was additionally transformed into *ATML1*/*atml1-1;pdf2-1* and Col-0. *Agrobacterium* strain GV3101 (MP90) was used for transformation and construction of transgenics by floral dip (Clough and Bent, 1998), followed by selection on 20 µg/ml hygromycin B. Segregation patterns of 3:1 for EYFP expression were observed among T2 progeny from at least 20 independent transformants, and representative homozygous T3 lines were selected for analysis. *Arabidopsis* plants were grown at 23°C under continuous light on soil comprised of Metro-Mix 380, vermiculite and perlite (4:3:2) (Hummert International). For RNA or lipid extraction, seeds were sterilized by chlorine gas treatment and sown onto 0.8% agar (Micropropagation Type II; Caisson Labs) containing Murashige and Skoog (MS) basal salts (Sigma-Aldrich) (Murashige and Skoog, 1962), 1% Suc, and 0.05% MES buffer at pH 5.8. Seeds were transferred to 23°C and grown under continuous light for 12 or 14 d. A razor blade was used to remove roots, and shoots were processed for RNA or lipid extraction. For growth under Pi limitation, vapor-sterilized seeds were germinated on Pi sufficient (P+) media (20.6 mM NH_4_NO_3_, 2.26 mM CaCl_2_ dihydrate, 0.759 mM MgSO_4_ heptahydrate, 18.8 mM KNO_3_ and 1.25 mM KH_2_PO_4_ monobasic, MS micronutrient solution (M529, PhytoTech Labs), 1% Suc, 0.05% MES, pH 5.7, 0.8% agar) or Pi limiting (P-) media (lacking KH_2_PO_4_ monobasic).

### Constructs for plant transformation

TAP constructs were generated by Gateway technology using *pKCTAP* and *pKNGSTAP* as described previously (Van Leene et al., 2011). PDF2 was amplified from an *Arabidopsis* cDNA library using PCR primers listed in **Supplemental Table 5**. The pdf2^ΔSTART^ constructs were generated by PCR amplification of PDF2 binary N’ and C’ TAP constructs using PCR primers flanking the START domain (**Supplemental Table 5**) followed by ligation. The SR54 binary vector for expression of GL2 under its native promoter (*proGL2:EYFP:GL2*) in plants was previously described (Schrick et al., 2014). To construct binary vectors expressing *PDF2* and *ATML1*, cDNA sequences were PCR amplified using Q5 High Fidelity Polymerase (New England Biolabs) and cloned into SR54 *proGL2:EYFP* cleaved with *Sal*I and *Kpn*I using NEBuilder HiFi DNA Assembly Master Mix (New England Biolabs) with gene-specific primers (**Supplemental Table 5**). The K107E and ΔSTART mutations in *PDF2* were generated using Q5 Site-Directed Mutagenesis Kit (New England Biolabs). The L480P mutation in *GL2* was generated by one-step PCR-based site-directed mutagenesis (Scott et al., 2002) using PfuUltra II Fusion HS DNA polymerase (Agilent Technologies) with primers listed in **Supplemental Table 5**. *HA:PDF2* and *HA:GL2* were constructed by transferring the respective cDNAs from pENTR/D-TOPO plasmids into pEarleyGate 201 (Earley et al., 2006) using Gateway LR Clonase II (Invitrogen).

### Tandem affinity purification

Affinity purification was performed as previously described (Luzarowski et al., 2017; Luzarowski et al., 2018). Whole cell native protein lysates (inputs) were harvested from *Arabidopsis* cell cultures expressing 35S:TAP:PDF2, 35S:PDF2:TAP, 35S:TAP:pdf2^ΔSTART^, 35S:pdf2^ΔSTART^:TAP, or empty vector. A soluble (membrane depleted) fraction was obtained by centrifugation of the lysate for 10 min at 14,000 rcf at 4°C, followed by ultracentrifugation for 1 h at 35,000 rcf at 4°C, and subsequent incubation with IgG Sepharose. After stringent washes, bait proteins were released from the beads by TEV protease cleavage. Samples were extracted as previously described (Giavalisco et al., 2011), using a methyl-tert-butyl ether (MTBE)/methanol/water solvent system to separate proteins, lipids, and polar compounds into pellet, organic, and aqueous phases, respectively. Following extraction, organic and aqueous phases were dried and stored at −20°C until LC/MS analysis.

### LC/MS analysis

Ultra-performance liquid chromatography (Waters Acquity UPLC System) coupled to an Exactive mass spectrometer (ThermoFisher Scientific) in positive and negative ionization mode was used to analyze the samples as described (Giavalisco et al., 2011). UPLC separation of the polar fraction was performed using an HSS T3 C18 reversed-phase column (100 mm × 2.1 mm × 1.8 μm particles; Waters). The mobile phases were 0.1% formic acid in water (Buffer A, ULC/MS; Biosolve) and 0.1% formic acid in acetonitrile (Buffer B, ULC/MS; Biosolve). A 2 μL sample (the dried-down aqueous fraction was resuspended in 200 μL of UPLC grade water) was loaded per injection. UPLC separation of the lipid fraction was performed using a C8 reversed-phase column (100 µm × 2.1 µm × 1.7 μm particles; Waters). Mobile phases were H_2_O (ULC/MS; Biosolve) with 1% 1 mM NH_4_Ac, 0.1% acetic acid (Buffer A) and acetonitrile:isopropanol (7:3, ULC/MS; Biosolve) containing 1% 1 mM NH_4_Ac, 0.1% acetic acid (Buffer B). A 2 μL sample (of the dried-down organic fraction resuspended in 200 μL of acetonitrile:isopropanol (7:3)) was loaded per injection. Processing of chromatograms, peak detection, and integration were performed using Refiner MS 12.0 (GeneData). Processing of mass spectrometry data included removal of isotopic peaks and of chemical noise, retention time alignment and adduct detection. Metabolic features (m/z at a given retention time) were queried against an in-house reference compound library (allowing 10 ppm error and up to 0.2 min deviation from the retention time).

### Recombinant protein production

The START domain coding region of PDF2 (PDF2(START)) was PCR amplified using gene-specific primers having ligation independent cloning (LIC) compatible extensions (**Supplemental Table 5**). Gel-purified PCR product and *Ssp*I-digested pET-His6-MBP-TEV-LIC vector (Addgene) were treated with T4 DNA polymerase with 25 mM dCTP and dGTP for 30 min at 22°C followed by heat inactivation. A 6 µl mixture of PCR product and vector was incubated at 22°C for 30 min, followed by addition of 1 µl 25 mM EDTA and *E. coli* transformation. Primers used to generate *pdf2^L467P^* via site-directed mutagenesis are listed in **Supplemental Table 5**. *E. coli* BL21 Rosetta 2 (DE3) (Novagen) cells carrying pET-His6-MBP-TEV-PDF2(START) and pdf2(START)^L467P^ were grown overnight in 5 mL LB with 40 µg/mL kanamycin at 37°C. The next day, 0.5 L freshly prepared media was inoculated with 1 mL of culture and growth was continued at 28°C. At OD_600_ 0.6, expression was induced with 0.5 mM IPTG (Sigma-Aldrich), followed by incubation at 16°C for 16 h. Cells were harvested by centrifugation at 4,000 rcf, 10 min at 4°C, the pellet was frozen in liquid nitrogen and stored at −20°C for 1 h. The cells were resuspended in 20 mL of ice-cold lysis buffer containing 50 mM sodium phosphate pH 7.4, 500 mM NaCl, 1 mM imidazole, 0.5 mM TCEP, 1 mM PMSF (Sigma-Aldrich), 10% glycerol, 0.1% [w/v] lysozyme (AppliChem) and cOmplete Protease Inhibitor Cocktail, EDTA free (Sigma-Aldrich). Bacterial slurry was sonicated in an ice-cold ultrasonic bath (RK 31, Bandelin) for 10 min, followed by centrifugation at 13,000 rcf for 10 min at 4°C. Supernatant was mixed with 2 mL of Ni-NTA agarose (Qiagen) on a rotary shaker for 1 h at 4°C. Ni-NTA beads with bound MBP-PDF2(START) protein were washed with 12 mL of ice-cold NaCl solutions. Protein was released from the beads using a step elution gradient (100-500 mM imidazole). Each step included 3 min incubations with 0.5 ml elution buffer containing 50 mM sodium phosphate pH 7.4, 500 mM NaCl, 0.5 mM TCEP, 1 mM PMSF, 10% glycerol, and increasing imidazole concentrations (100-500 mM). Concentration and purity of MBP-PDF2(START) in elution fractions was estimated by SDS-PAGE. Imidazole was removed and proteins were concentrated using Amicon Ultra 15 mL centrifugal filters having 10 kDa cut-off. Protein folding was assessed using nano differential scanning fluorimetry (nanoDSF). Aliquots of purified protein were stored at −20°C in 50 mM sodium phosphate buffer (pH 7.4) supplemented with 500 mM NaCl.

### Liposome preparation

Lipids (Avanti Polar Lipids (Alabaster, AL) were dissolved in chloroform. A total of 5 mg lipid for each liposome batch was dried in a glass tube under N_2_ at 60°C. Residual chloroform was removed under vacuum overnight. Dried lipid cakes were rehydrated in 500 mM NaCl and 50 mM sodium phosphate (pH 7.4) at room temperature. Small unilamellar vesicles (SUVs) were formed by sonication using an ultrasonic bath (RK 31, Bandelin) for 15 min or by extrusion through two layers of polycarbonate membranes with 50 nm pore size (Nuclepore hydrophilic membrane, Whatman) in a handheld extruder (Avanti Polar Lipids) or by sonication. Hydrodynamic radii of liposomes were determined by dynamic light scattering (DLS) to validate successful SUV formation.

### Microscale thermophoresis (MST)

MST measurements were performed using a Monolith NT.115 (NanoTemper). Capillaries were loaded into the instrument assets in 16-point ligand titrations. MBP-PDF2(START), MBP-PDF2(START)^L467P^ were labeled in 50 mM sodium phosphate buffer (pH 7.4) supplemented with 500 mM NaCl using Monolith Protein Labeling kit RED-MALEIMIDE (NanoTemper) according to manufacturer’s instructions. To remove the MBP tag, labeled proteins were incubated with Ni-NTA agarose (Qiagen) on a rotary shaker for 1 h at RT. Ni-NTA beads were washed with 50 mM sodium phosphate buffer (pH 7.4) supplemented with 500 mM NaCl prior to release with two rounds of TEV protease digestion, each with 30 U of TEV for 1 h at RT. Binding was performed in 50 mM sodium phosphate buffer (pH 7.4) supplemented with 500 mM NaCl using standard capillaries. MO. Affinity Analysis software (NanoTemper) was used to analyze binding affinities from changes in fluorescence. SDS-Test was performed according to the NanoTemper MST manual to exclude that observed changes in fluorescence were due to ligand induced changes in protein aggregation.

### RNA extraction and quantitative real-time PCR

Plant samples of ∼50 mg were frozen in liquid nitrogen and stored at −80°C prior to RNA extraction with RNeasy Plant Mini Kit and on-column RNase-Free DNase Set (Qiagen). Total RNA (0.5 µg) was used as a template for cDNA synthesis with GoScript Reverse Transcriptase (Promega). qRT-PCR was performed using iTaq SYBR Green Supermix with the CFX96 Touch Real-Time PCR Detection System (Bio-Rad) with gene-specific primers (**Supplemental Table 5**). Reactions contained 10 μL SYBR Green Supermix, 1 μL forward and reverse 10 µM primers, and 5 μL cDNA (diluted 5-fold) in 20 µL. Standard curves were generated from 10-fold dilutions of amplicons for each primer pair. *ACT7* served as the reference gene. Data represent four biological samples of seedling shoots with three technical replicates for each biological sample.

### Lipid extraction from plant material

Plant tissues were transferred to hot isopropanol (70°C) with 0.01% BHT (butylated hydroxytoluene, Sigma) for 15 min followed by cooling to room temperature, and storage at −80°C prior to processing. Lipid extraction was done with chloroform: (isopronanol + methanol):water (30:65:3.5). Samples were incubated overnight at 50-100 rpm at room temperature followed by solvent evaporation. Extracted lipids were transferred to 2 ml glass vials and dried under N_2_. Based on lipid dry weight and formula weight of ∼800 Da, lipids were eluted at 100 mM with chloroform. 100 µl of a 100 µM lipid mixture was dried under N_2_ and stored at −80°C prior to analysis. Dried lipid fractions were resuspended in 200 μL UPLC-grade acetonitrile:isopropanol (7:3). A 2 μL sample was loaded per injection. LC/MS analysis was performed as described above. Raw intensities were normalized to the median of chromatogram intensity.

### Imaging of plants and quantification of trichomes and roots

Seedlings, trichome phenotypes, and EYFP expression were imaged with a Leica M125 fluorescence stereo microscope fitted with a GFP2 filter set, a Leica DFC295 digital camera with Leica Application Suite 4.1. Trichome quantification was performed as previously described (Schrick et al., 2014). Root lengths were measured using ImageJ software analysis of seedling images from BioRad Gel Doc XR+ Imaging System. Mature plants were imaged with a Canon PowerShot ELPH 350 HS digital camera.

### *In vitro* transcription and translation and electrophoretic mobility shift assay (EMSA)

*PDF2* and *ATML1* cDNAs were cloned from pENTR/SD/D-TOPO vectors (ABRC) into pIX-HALO (ABRC) using Gateway LR Clonase II Enzyme mix (ThermoFisher Scientific). The K107E mutation in *PDF2* was generated using Q5 Site-Directed Mutagenesis Kit (New England Biolabs) with described primers (**Supplemental Table 5**). Halo fusion proteins were produced from 1.5 µg plasmid DNA in a 15 µL reaction using TNT SP6 High-Yield Wheat Germ Protein Expression System (Promega). Protein expression was confirmed by Western blot with Anti-HaloTag monoclonal Ab (1:2000) (Promega). Cy3-labeled and unlabeled dsDNA probes were generated with oligonucleotides listed in **Supplemental Table 5**. Annealing was performed with 25 µM oligonucleotides in 100 mM Tris-Cl (pH 7.5), 1 M NaCl, 10 mM EDTA at 95°C for 2 min, followed by 57°C for 5 min, 37°C for 90 min and 37°C for 2 min. EMSA reactions (20 µL) were prepared as previously described (Evens et al., 2017), with the following modifications: 6 µl of *in vitro* translated product was pre-incubated with binding buffer at 28°C for 10 min. Binding reactions were initiated by adding 200 nM of Cy3-labeled probe, followed by a 20 min incubation. After electrophoresis in a 0.6 % agarose gel (1X TBE, pH 8.3) at 150 V for 1 h at 4°C, the protein-DNA complexes were analysed with a Typhoon Trio Imager (GE Healthcare) using the 532 nm laser and 580 nm emission filter.

### *In vivo* protein stability assay

At 5-6 days after germination on P+ or P-agar media, 20-30 seedlings per sample were transferred to liquid P+ or P-media and growth was continued for 16 h at 23°C under continuous light. Cycloheximide (Sigma-Aldrich) (400 µM final concentration) or DMSO was added at 0 h, and harvesting occurred at 0, 2, 5, 10 or 24 h. For proteasome inhibition experiments, cycloheximide was added together with MG132 (50 µM) (Sigma Aldrich) or DMSO control at 0 h. Seedling samples were frozen in liquid nitrogen and stored at −80°C prior to protein extraction. Tissue was homogenized in liquid nitrogen and hot SDS buffer (8 M urea, 2% SDS, 0.1 M DTT, 20% glycerol, 0.1 M Tris pH 6.8, 0.004% bromophenol blue) was added prior to SDS-PAGE and Western blotting. Anti-HA (1:10,000; Pierce) or Anti-GFP (1:2000; Roche) served as primary Abs, followed by Goat Anti-Mouse IgG [HRP] (1:3000; GenScript A00160) as the secondary. Proteins were detected with SuperSignal West Femto Maximum Sensitivity Substrate (ThermoFisher Scientific) using Azure 300 chemiluminescence imager (Azure Biosystems), and blots were stained with Bio-Safe Coomassie Blue G-250 (Bio-Rad) to monitor protein loading. Band intensities were quantified with ImageJ.

### Statistical analysis

#### TAP experiments

GeneData derived log_2_ transformed raw metabolite intensities were used for statistical analysis (**Supplemental Data Set 1)**. MaxQuant derived log_2_ transformed LFQ protein intensities were used for statistical analysis. Unpaired *t*-test (two-tailed distribution) was used to assess significance. Data are from six (soluble fraction; experiment 2) samples. Samples represent individual pull-downs, performed in parallel, using same starting material harvested from two independent lines tagged on either amino or carboxyl terminus.

#### Quantitative real-time PCR

The data represent four biological samples of seedling shoots from wild type and each mutant, with three technical replicates for each biological sample. Standard curves were generated for each primer pair and used to calculate relative units for the experimental samples. All gene expression data were normalized to *ACT7* (AT5G09810) as the reference gene. Unpaired *t*-tests (two-tailed distribution) were used to assess significance (p < 0.05) for gene expression differences between wild type and mutant.

#### Lipidomics

GeneData derived raw lipid intensities were normalized to the median intensity of all mass features detected in a given chromatogram and used for statistical analysis. The data represent 4-5 biological replicates of seedling shoots from wild type and each of the mutants. For each lipid, averages and standard deviations of lipid intensities in the 4-5 replicates were determined for the wild-type and mutant samples (**Supplemental Data Sets 3 and 4**). The fold-changes (mutant/wild-type) were calculated for each lipid. One-way Anova (Tukey’s test) was used for comparison of lipid classes for > 3 genotypes. Unpaired *t*-test with two-tailed distribution was used to test significance of changes greater than 2-fold (p ≤ 0.05). Critical data sets were subjected to multiple testing for significance in MetaboAnalyst (Xia et al., 2009), correcting for FDR (**Supplemental Tables 1-4**).

#### Protein stability assays

The graphed data represent three or four independent cycloheximide experiments with whole seedlings. Numerical values for protein levels were obtained from band intensity quantification and were normalized to the mock zero (M0) samples, which were designated a value of 1.0. Protein half-life (in h) was determined by the protein level at a *y*-axis value of 0.5. Unpaired *t*-tests (two-tailed distribution) were used to assess significance (p < 0.05) for protein level differences between P+ and P-samples at each time point.

### Accession numbers

*Arabidopsis thaliana* HD-Zip IV TFs: PDF2 (At4g04890), GL2 (At1g79840), ATML1 (At4g21750); *Physcomitrium patens* HD-Zip IV TF: PpHDZIV (XP_024401280.1), *Homo sapiens* START domain proteins: StAR/STARD1 (NP_000340.2); *Homo sapiens* PCTP/STARD2 (NP_067036.2); *Arabidopsis thaliana* phospholipid catabolism enzymes: GDPD1 (At3g02040), GDPD2 (At5g41080), GDPD3 (At5g43300), NPC2 (At2g26870), NPC4 (At3g03530), NPC6 (At3g48610), PLDɛ (At1g55180), PLDζ1 (At3g16785), PLDζ2 (At3g05630).

## Supporting information

Supplemental Data Sets

## Supplemental Data

**Supplemental Figure 1.** Lysophosphatidylcholines bind to the START domain of PDF2 *in vitro*.

**Supplemental Figure 2.** The *atml1-1;pdf2-1* double mutant exhibits severely altered lipid composition.

**Supplemental Figure 3.** Heat maps illustrate lipidomic profiles in wild type, *pdf2*, *atml1* and *gl2* seedling shoots under normal growth conditions.

**Supplemental Figure 4.** Heat maps illustrate lipidomic profiles in wild type, *pdf2*, *atml1* and *gl2* mutants under Pi sufficient and limiting conditions.

**Supplemental Figure 5.** Volcano plots reveal lipid changes in HD-Zip IV mutants under Pi sufficient and limiting conditions.

**Supplemental Figure 6.** Plant phenotypes from START-domain dependent expression of PDF2.

**Supplemental Figure 7.** Protein stability of PDF2 and GL2 is reduced under Pi limitation and START domain mutant exhibits enhanced protein instability.

**Supplemental Table 1.** Lipidomic changes in *atml1;pdf2-1* vs. wild type.

**Supplemental Table 2.** Lipidomic changes in *pdf2-1* vs. wild type under Pi limitation.

**Supplemental Table 3.** Lipidomic changes in *pdf2-2* vs. wild type under Pi limitation.

**Supplemental Table 4.** Lipidomic changes in *EYFP:pdf2*^ΔSTART^ vs. *EYFP:PDF2* under Pi limitation.

**Supplemental Table 5.** Oligonucleotides used in this study.

**Supplemental Data Sets 1A and 1B.** TAP lipidomics data for the PDF2 TF from soluble fractions.

**Supplemental Data Set 2A.** List of putative transcriptional target genes from DAP-seq data for PDF2.

**Supplemental Data Set 2B.** GO enrichment for DAP-seq targets of PDF2.

**Supplemental Data Set 3A.** Comprehensive lipidomic data from wild-type and *pdf2*, *atml1;pdf2*, *atml1*, and *gl2* mutants.

**Supplemental Data Set 3B.** Maximum normalized comprehensive lipidomic data from wild-type and *pdf2*, *atml1;pdf2*, *atml1*, and *gl2* mutants.

**Supplemental Data Set 4A.** Lipidomic data from wild-type, *pdf2*, *atml1*, and *gl2* mutants, as well as *EYFP:PDF2*, *EYFP:pdf2*^Δ^*^ST^*, *EYFP:GL2*, and *EYFP:gl2^L480P^* transgenic lines in Pi sufficient (P+) and Pi limiting (P-) media.

**Supplemental Data Set 4B.** Maximum normalized lipidomic data from wild-type, *pdf2*, *atml1*, and *gl2* mutants, as well as *EYFP:PDF2*, *EYFP:pdf2^ΔST^*, *EYFP:GL2*, and *EYFP:gl2^L480P^* transgenic lines in Pi sufficient (P+) and Pi limiting (P-) media.

**Supplemental Data Set 4C.** Volcano plot calculations for lipidomic data from wild-type, *pdf2*, *atml1*, and *gl2* mutants, as well as *EYFP:PDF2*, *EYFP: pdf2^ΔST^*, *EYFP:GL2*, and *EYFP: gl2^L480P^* transgenic lines in Pi sufficient (P+) and Pi limiting (P-) media.

**Supplemental Data Set 5A.** Lipidomics Mass Spectrometry Details: All Detected Peaks, Putative Metabolite Name Identified: Tandem Affinity Purification Experiment.

**Supplmental Data Set 5B.** Lipidomics Mass Spectrometry Details: All Detected Peaks, Putative Metabolite Name Identified: Mutants Experiment.

**Supplemental Data Set 5C.** Lipidomics Mass Spectrometry Details: All Detected Peaks, Putative Metabolite Name Identified: Pi Limitation Experiment.

## ACKNOWLEDGMENTS

This research was funded by the National Science Foundation (MCB1616818), National Institute of General Medical Sciences of the National Institute of Health under Award no. P20GM103418, USDA National Institute of Food and Agriculture Hatch/Multi-State project 1013013, and Johnson Cancer Research Center at Kansas State University. This is contribution no. 20-003-J from the Kansas Agricultural Experiment Station. We thank Anne Michaelis for processing lipidomics samples, Mary Roth for help with lipid extraction, and Adrienne Roeder, Xuemin Wang, Ruth Welti and Lothar Willmitzer for valuable input.

## AUTHOR CONTRIBUTIONS

I.W., A.S. and K.S conceived of the experiments and wrote the manuscript. X.H. and A.K. designed and developed constructs for recombinant expression in *E. coli*. K.S. and G.L.M. performed RNA extractions and qRT-PCR and G.L.M analyzed DAP-seq data. I.W. and J.S. performed TAP experiments and analyzed data. I.W. and J.G. purified recombinant protein and A.S. performed liposome binding experiments. K.S. prepared protein alignment and structural models. P.K.-B., A.T. and D.K.H. prepared liposomes. K.S., A.K., T.M., K.A.T. and S.T.P designed and constructed plasmids for DNA binding and plant assays. T.M. and K.S. performed lipid extractions, DNA binding and cycloheximide assays. S.T.P. and K.S. conducted root growth assays and EYFP expression analysis, and K.S. performed trichome quantification.

## REFERENCES

Abe, M., Katsumata, H., Komeda, Y., and Takahashi, T. (2003). Regulation of shoot epidermal cell differentiation by a pair of homeodomain proteins in Arabidopsis. Development 130, 635–643.

Alpy, F., and Tomasetto, C. (2005). Give lipids a START: the StAR-related lipid transfer (START) domain in mammals. J.Cell Sci. 118, 2791–2801.

Alpy, F., Legueux, F., Bianchetti, L., and Tomasetto, C. (2009). [START domain-containing proteins: a review of their role in lipid transport and exchange]. Med Sci (Paris) 25, 181–191.

Bose, H.S., Sugawara, T., Strauss, J.F., 3rd, Miller, W.L., and International Congenital Lipoid Adrenal Hyperplasia, C. (1996). The pathophysiology and genetics of congenital lipoid adrenal hyperplasia. The New England journal of medicine 335, 1870–1878.

Cheng, Y., Zhou, W., El Sheery, N.I., Peters, C., Li, M., Wang, X., and Huang, J. (2011). Characterization of the Arabidopsis glycerophosphodiester phosphodiesterase (GDPD) family reveals a role of the plastid-localized AtGDPD1 in maintaining cellular phosphate homeostasis under phosphate starvation. Plant J 66, 781–795.

Clark, B.J. (2020). The START-domain proteins in intracellular lipid transport and beyond. Mol Cell Endocrinol 504, 110704.

Clough, S.J., and Bent, A.F. (1998). Floral dip: a simplified method for Agrobacterium-mediated transformation of Arabidopsis thaliana. Plant J 16, 735–743.

Drissner, D., Kunze, G., Callewaert, N., Gehrig, P., Tamasloukht, M., Boller, T., Felix, G., Amrhein, N., and Bucher, M. (2007). Lyso-phosphatidylcholine is a signal in the arbuscular mycorrhizal symbiosis. Science 318, 265–268.

Earley, K.W., Haag, J.R., Pontes, O., Opper, K., Juehne, T., Song, K., and Pikaard, C.S. (2006). Gateway-compatible vectors for plant functional genomics and proteomics. Plant J 45, 616–629.

Evens, N.P., Buchner, P., Williams, L.E., and Hawkesford, M.J. (2017). The role of ZIP transporters and group F bZIP transcription factors in the Zn-deficiency response of wheat (Triticum aestivum). Plant J 92, 291–304.

Fluck, C.E., Maret, A., Mallet, D., Portrat-Doyen, S., Achermann, J.C., Leheup, B., Theintz, G.E., Mullis, P.E., and Morel, Y. (2005). A novel mutation L260P of the steroidogenic acute regulatory protein gene in three unrelated patients of Swiss ancestry with congenital lipoid adrenal hyperplasia. J Clin Endocrinol Metab 90, 5304–5308.

Giavalisco, P., Li, Y., Matthes, A., Eckhardt, A., Hubberten, H.M., Hesse, H., Segu, S., Hummel, J., Kohl, K., and Willmitzer, L. (2011). Elemental formula annotation of polar and lipophilic metabolites using (13) C, (15) N and (34) S isotope labelling, in combination with high-resolution mass spectrometry. Plant J 68, 364–376.

Gou, J.Y., Li, K., Wu, K., Wang, X., Lin, H., Cantu, D., Uauy, C., Dobon-Alonso, A., Midorikawa, T., Inoue, K., Sanchez, J., Fu, D., Blechl, A., Wallington, E., Fahima, T., Meeta, M., Epstein, L., and Dubcovsky, J. (2015). Wheat Stripe Rust Resistance Protein WKS1 Reduces the Ability of the Thylakoid-Associated Ascorbate Peroxidase to Detoxify Reactive Oxygen Species. Plant Cell 27, 1755–1770.

Hirashima, T., Jimbo, H., Kobayashi, K., and Wada, H. (2021). A START domain-containing protein is involved in the incorporation of ER-derived fatty acids into chloroplast glycolipids in Marchantia polymorpha. Biochem Biophys Res Commun 534, 436–441.

Hong, Y., Devaiah, S.P., Bahn, S.C., Thamasandra, B.N., Li, M., Welti, R., and Wang, X. (2009). Phospholipase D epsilon and phosphatidic acid enhance Arabidopsis nitrogen signaling and growth. Plant J 58, 376–387.

Kamata, N., Okada, H., Komeda, Y., and Takahashi, T. (2013a). Mutations in epidermis-specific HD-ZIP IV genes affect floral organ identity in Arabidopsis thaliana. Plant J.

Kamata, N., Sugihara, A., Komeda, Y., and Takahashi, T. (2013b). Allele-specific effects of PDF2 on floral morphology in Arabidopsis thaliana. Plant Signal Behav 8, e27417.

Kanno, K., Wu, M.K., Agate, D.S., Fanelli, B.J., Wagle, N., Scapa, E.F., Ukomadu, C., and Cohen, D.E. (2007). Interacting proteins dictate function of the minimal START domain phosphatidylcholine transfer protein/StarD2. J.Biol.Chem. 282, 30728–30736.

Khosla, A., Paper, J.M., Boehler, A.P., Bradley, A.M., Neumann, T.R., and Schrick, K. (2014). HD-Zip Proteins GL2 and HDG11 Have Redundant Functions in Arabidopsis Trichomes, and GL2 Activates a Positive Feedback Loop via MYB23. Plant Cell 26, 2184–2200.

Letourneau, D., Lefebvre, A., Lavigne, P., and LeHoux, J.G. (2015). The binding site specificity of STARD4 subfamily: Breaking the cholesterol paradigm. Mol Cell Endocrinol 408, 53–61.

Letourneau, D., Lorin, A., Lefebvre, A., Frappier, V., Gaudreault, F., Najmanovich, R., Lavigne, P., and LeHoux, J.G. (2012). StAR-related lipid transfer domain protein 5 binds primary bile acids. Journal of Lipid Research 53, 2677–2689.

Li, M., Welti, R., and Wang, X. (2006). Quantitative profiling of Arabidopsis polar glycerolipids in response to phosphorus starvation. Roles of phospholipases D zeta1 and D zeta2 in phosphatidylcholine hydrolysis and digalactosyldiacylglycerol accumulation in phosphorus-starved plants. Plant Physiol 142, 750–761.

Lin, W.D., Liao, Y.Y., Yang, T.J., Pan, C.Y., Buckhout, T.J., and Schmidt, W. (2011). Coexpression-based clustering of Arabidopsis root genes predicts functional modules in early phosphate deficiency signaling. Plant Physiol 155, 1383–1402.

Luzarowski, M., Wojciechowska, I., and Skirycz, A. (2018). 2 in 1: One-step Affinity Purification for the Parallel Analysis of Protein-Protein and Protein-Metabolite Complexes. J Vis Exp, 57720.

Luzarowski, M., Kosmacz, M., Sokolowska, E., Jasinska, W., Willmitzer, L., Veyel, D., and Skirycz, A. (2017). Affinity purification with metabolomic and proteomic analysis unravels diverse roles of nucleoside diphosphate kinases. J Exp Bot 68, 3487–3499.

Mi, H.Y., Muruganujan, A., Ebert, D., Huang, X.S., and Thomas, P.D. (2019). PANTHER version 14: more genomes, a new PANTHER GO-slim and improvements in enrichment analysis tools. Nucleic Acids Res 47, D419–D426.

Mongrand, S., Cassagne, C., and Bessoule, J.J. (2000). Import of lyso-phosphatidylcholine into chloroplasts likely at the origin of eukaryotic plastidial lipids. Plant Physiol 122, 845–852.

Murashige, T., and Skoog, F. (1962). A Revised Medium for Rapid Growth and Bio Assays with Tobacco Tissue Cultures. Physiologia Plantarum 15, 473–497.

Nagata, K., Ishikawa, T., Kawai-Yamada, M., Takahashi, T., and Abe, M. (2021). Ceramides mediate positional signals in Arabidopsis thaliana protoderm differentiation. Development 148.

Nakamura, Y., Awai, K., Masuda, T., Yoshioka, Y., Takamiya, K., and Ohta, H. (2005). A novel phosphatidylcholine-hydrolyzing phospholipase C induced by phosphate starvation in Arabidopsis. J Biol Chem 280, 7469–7476.

Nobusawa, T., Okushima, Y., Nagata, N., Kojima, M., Sakakibara, H., and Umeda, M. (2013). Synthesis of very-long-chain fatty acids in the epidermis controls plant organ growth by restricting cell proliferation. PLoS Biol 11, e1001531.

O’Malley, R.C., Huang, S.C., Song, L., Lewsey, M.G., Bartlett, A., Nery, J.R., Galli, M., Gallavotti, A., and Ecker, J.R. (2016). Cistrome and Epicistrome Features Shape the Regulatory DNA Landscape. Cell 166, 1598.

Ogawa, E., Yamada, Y., Sezaki, N., Kosaka, S., Kondo, H., Kamata, N., Abe, M., Komeda, Y., and Takahashi, T. (2015). ATML1 and PDF2 Play a Redundant and Essential Role in Arabidopsis Embryo Development. Plant Cell Physiol 56, 1183–1192.

Ohashi, Y., Oka, A., Rodrigues-Pousada, R., Possenti, M., Ruberti, I., Morelli, G., and Aoyama, T. (2003). Modulation of phospholipid signaling by GLABRA2 in root-hair pattern formation. Science 300, 1427–1430.

Pant, B.D., Burgos, A., Pant, P., Cuadros-Inostroza, A., Willmitzer, L., and Scheible, W.R. (2015). The transcription factor PHR1 regulates lipid remodeling and triacylglycerol accumulation in Arabidopsis thaliana during phosphorus starvation. J Exp Bot 66, 1907–1918.

Peterson, K.M., Shyu, C., Burr, C.A., Horst, R.J., Kanaoka, M.M., Omae, M., Sato, Y., and Torii, K.U. (2013). Arabidopsis homeodomain-leucine zipper IV proteins promote stomatal development and ectopically induce stomata beyond the epidermis. Development 140, 1924–1935.

Ponting, C.P., and Aravind, L. (1999). START: a lipid-binding domain in StAR, HD-ZIP and signalling proteins. Trends Biochem.Sci. 24, 130–132.

Roderick, S.L., Chan, W.W., Agate, D.S., Olsen, L.R., Vetting, M.W., Rajashankar, K.R., and Cohen, D.E. (2002). Structure of human phosphatidylcholine transfer protein in complex with its ligand. Nat.Struct.Biol. 9, 507–511.

Roeder, A.H., Cunha, A., Ohno, C.K., and Meyerowitz, E.M. (2012). Cell cycle regulates cell type in the Arabidopsis sepal. Development 139, 4416–4427.

Rombola-Caldentey, B., Rueda-Romero, P., Iglesias-Fernandez, R., Carbonero, P., and Onate-Sanchez, L. (2014). Arabidopsis DELLA and two HD-ZIP transcription factors regulate GA signaling in the epidermis through the L1 box cis-element. Plant Cell 26, 2905–2919.

Roy, A., Kucukural, A., and Zhang, Y. (2010). I-TASSER: a unified platform for automated protein structure and function prediction. Nat Protoc 5, 725–738.

Rubio, V., Linhares, F., Solano, R., Martin, A.C., Iglesias, J., Leyva, A., and Paz-Ares, J. (2001). A conserved MYB transcription factor involved in phosphate starvation signaling both in vascular plants and in unicellular algae. Genes Dev 15, 2122–2133.

Savaldi-Goldstein, S., Peto, C., and Chory, J. (2007). The epidermis both drives and restricts plant shoot growth. Nature 446, 199–202.

Schrick, K., Nguyen, D., Karlowski, W.M., and Mayer, K.F. (2004). START lipid/sterol-binding domains are amplified in plants and are predominantly associated with homeodomain transcription factors. Genome Biol 5, R41.

Schrick, K., Bruno, M., Khosla, A., Cox, P.N., Marlatt, S.A., Roque, R.A., Nguyen, H.C., He, C., Snyder, M.P., Singh, D., and Yadav, G. (2014). Shared functions of plant and mammalian StAR-related lipid transfer (START) domains in modulating transcription factor activity. BMC Biol 12, 70.

Scott, S.P., Teh, A., Peng, C., and Lavin, M.F. (2002). One-step site-directed mutagenesis of ATM cDNA in large (20kb) plasmid constructs. Hum Mutat 20, 323.

Sluchanko, N.N., Tugaeva, K.V., Faletrov, Y.V., and Levitsky, D.I. (2016). High-yield soluble expression, purification and characterization of human steroidogenic acute regulatory protein (StAR) fused to a cleavable Maltose-Binding Protein (MBP). Protein expression and purification 119, 27–35.

Su, Y., Li, M., Guo, L., and Wang, X. (2018). Different effects of phospholipase Dzeta2 and non-specific phospholipase C4 on lipid remodeling and root hair growth in Arabidopsis response to phosphate deficiency. Plant J 94, 315–326.

Takada, S., Takada, N., and Yoshida, A. (2013). ATML1 promotes epidermal cell differentiation in Arabidopsis shoots. Development 140, 1919–1923.

Tillman, M.C., Imai, N., Li, Y., Khadka, M., Okafor, C.D., Juneja, P., Adhiyaman, A., Hagen, S.J., Cohen, D.E., and Ortlund, E.A. (2020). Allosteric regulation of thioesterase superfamily member 1 by lipid sensor domain binding fatty acids and lysophosphatidylcholine. Proc Natl Acad Sci U S A 117, 22080–22089.

Van Leene, J., Eeckhout, D., Persiau, G., Van De Slijke, E., Geerinck, J., Van Isterdael, G., Witters, E., and De Jaeger, G. (2011). Isolation of transcription factor complexes from Arabidopsis cell suspension cultures by tandem affinity purification. Methods Mol Biol 754, 195–218.

Xia, J., Psychogios, N., Young, N., and Wishart, D.S. (2009). MetaboAnalyst: a web server for metabolomic data analysis and interpretation. Nucleic Acids Res 37, W652–660.

Yang, B., Li, M., Phillips, A., Li, L., Ali, U., Li, Q., Lu, S., Hong, Y., Wang, X., Guo, L. (2021). Nonspecific phospholipase C4 hydrolyzes phosphosphingolipids and sustains plant root growth during phosphate deficiency. Plant Cell, 1–15.

Yang, J., and Zhang, Y. (2015). I-TASSER server: new development for protein structure and function predictions. Nucleic Acids Res 43, W174–181.

