## Supplemental Figures and Tables for "*Arabidopsis* PROTODERMAL FACTOR2 binds lysophosphatidylcholines and transcriptionally regulates phospholipid metabolism"

[†] submission posthumously

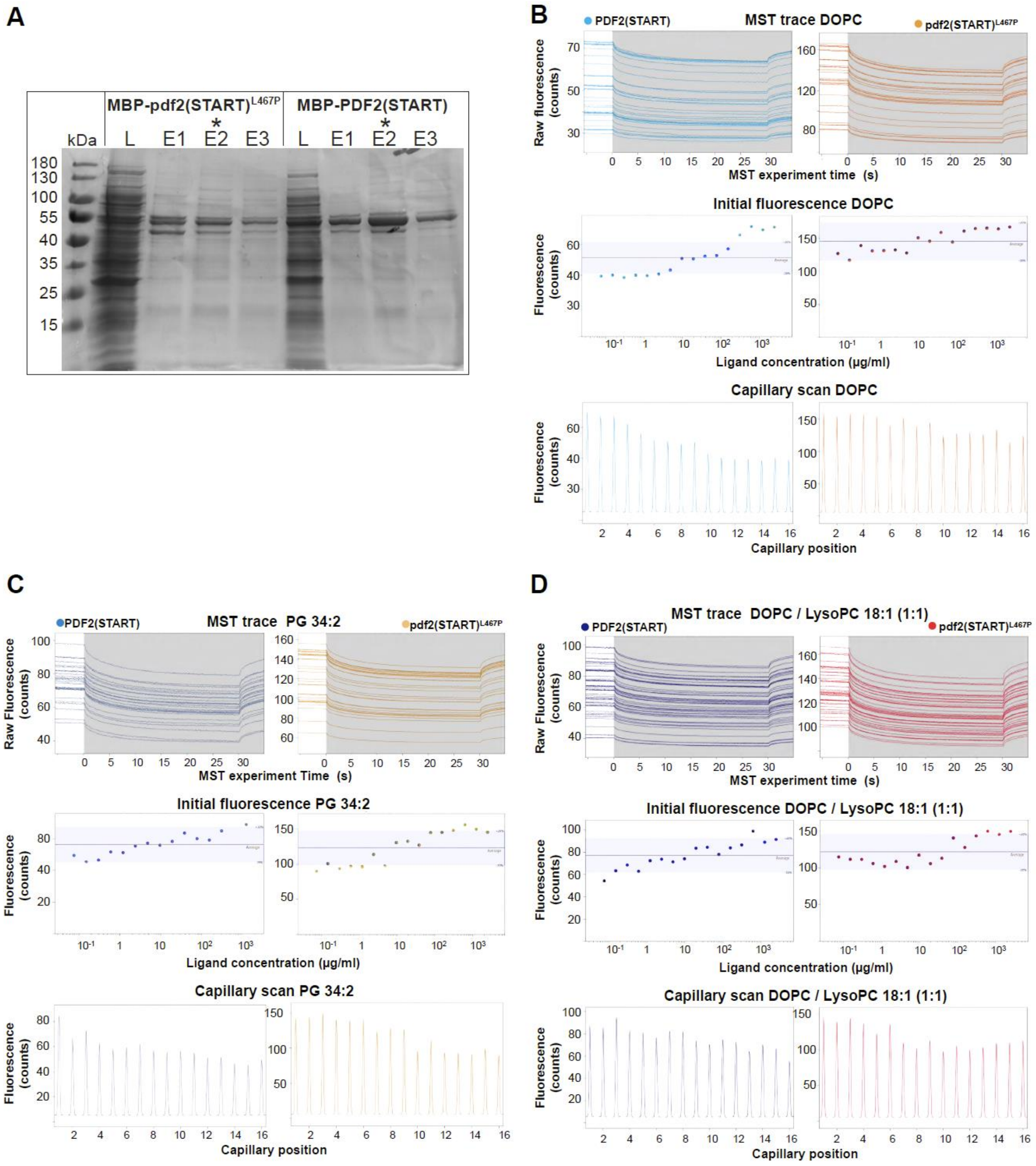


**Supplemental Figure 1.** Lysophosphatidylcholines bind to the START domain of PDF2 *in vitro*. Related to **Figure 2F**

**(A)** Purification of PDF2(START) mutant and wild-type proteins used in the liposome binding experiments. Coomassie blue-stained gel shows samples from the purification of maltose binding protein (MBP = His6-MBP-TEV) tagged mutant pdf2(START)$^{L467P}$ and wild-type PDF2(START). L, Lysate; E1, E2, and E3, Elutions with 100, 250, and 500 mM imidazole, respectively. Asterisks indicate samples that were used for the liposome binding experiments followed by microscale thermophoresis (MST). Note that the His6-MBP tag was removed by TEV protease cleavage prior to binding analysis.

**(B-D)** Exemplary microscale thermophoresis (MST) traces (top). Binding to liposomes results in changes in PDF2(START) and pdf2(START)$^{L467P}$ fluorescence used to determine binding and binding affinities. Fluorescence increases upon liposome binding, reminiscent of protein unfolding. Initial fluorescence across ligand titrations (middle). Capillary scans across ligand titrations (bottom). Capillary 1 corresponds to the highest and capillary 16 to the lowest ligand concentration. **(B)** DOPC (PC (36:2)) liposomes in MST fluorescence measurements. **(C)** PG 34:2 liposomes in MST fluorescence measurements. **(D)** DOPC / lysophosphatidylcholine (LysoPC) 18:1 liposomes in MST fluorescence measurements.

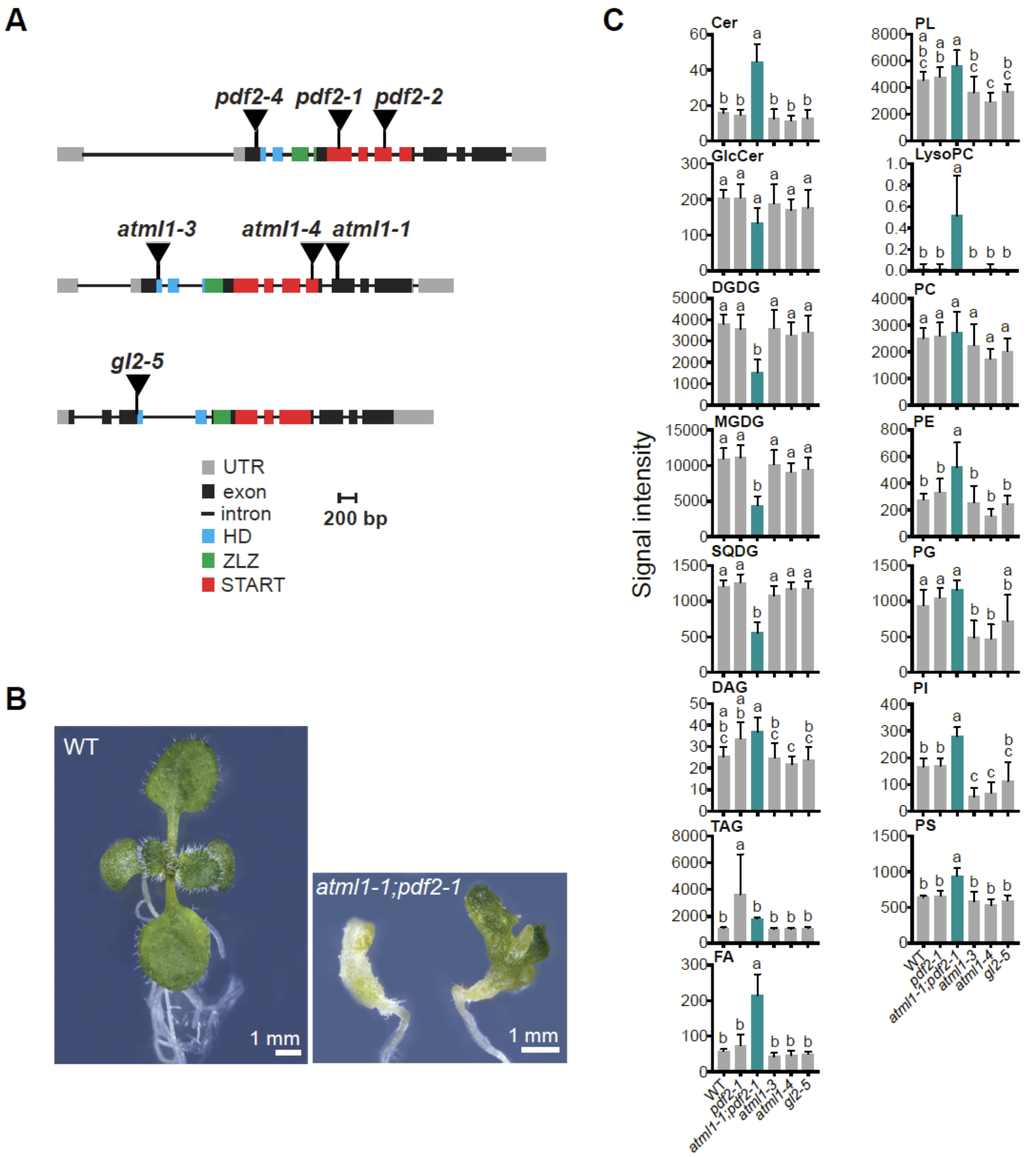


**Supplemental Figure 2.** The *atml1-1;pdf2-1* double mutant exhibits severely altered lipid composition. Related to **Supplemental Data Set 3A.**

**(A)** Map positions of T-DNA insertions for *PDF2*, *ATML1* and *GL2*. Schematic of genomic regions encoding *PDF2*, *ATML1* and *GL2*. T-DNA insertions are shown relative to the gene segments encoding the Homeodomain (HD), Zipper Loop Zipper (ZLZ), and START domain.

**(B)** Representative 12-d-old seedlings from wild type (WT) and *atml1-1;pdf2-1* double mutant. Size bars = 1.0 mm.

**(C)** Lipids were extracted from 12-d-old seedling shoots from WT, *pdf2-1*, *atml1-1;pdf2-1*, *atml1-3*, *atml1-4*, and *gl2-5*, and processed for lipidomics by LC-MS. Lipid profiles indicate relative intensities of ceramides (Cer), glucosylceramides (GlcCer), digalactosyldiacylglycerols (DGDG), monogalactosyldiacylglycerols (MGDG), sulfoquinovosyl diacylglycerols (SQDG), diacylglyerols (DAG), triacylglycerols (TAG), and fatty acids (FA) (left) and total phospholipids (PL), phosphatidylcholine (PC), phosphatidylethanolamine (PE), phosphatidylglycerol (PG), phosphoinositol (PI), and phosphatidylserine (PS) (right). Error bars indicate SD for n = 5 biological replicates. Significant differences between genotypes determined by one-way ANOVA, Tukey's test, and indicated by letters: $p < 0.05$.

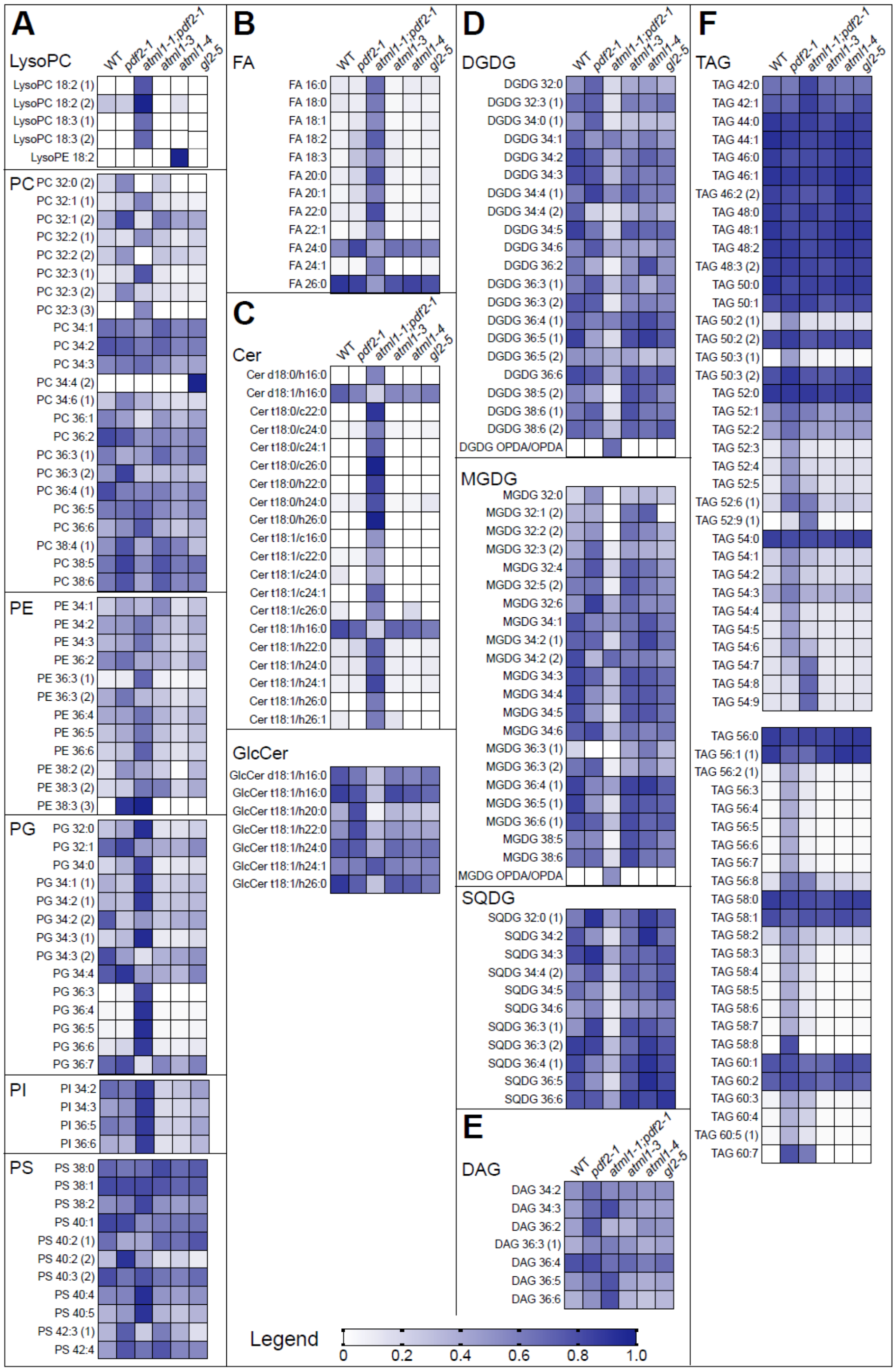


**Supplemental Figure 3.** Heat maps illustrate lipidomic profiles in wild type, *pdf2*, *atml1* and *gl2* seedling shoots under normal growth conditions. Related to **Supplemental Figure 2** and **Supplemental Table 3B.**

**(A-F)** Lipidomics data is shown for 12-d-old seedlings whose roots were excised from wild-type (WT), *pdf2-1*, *atml1-1;pdf2-1*, *atml1-3*, *atml1-4* and *gl2-5* genotypes. Minimum and maximum values were normalized to 0.0 and 1.0, respectively, for visualization purposes. **(A)** Phospholipids (LysoPC, PC, PE, PG, PI, PS), **(B)** Fatty acids (FA), **(C)** Ceramides (Cer) and glucosylceramides (GlcCer), **(D)** Digalactosyldiacylglycerols (DGDG), monogalactosyldiacylglycerols (MGDG) and sulfoquinovosyl diacylglycerols (SQDG), **(E)** Diacylglycerols (DAG), and **(F)** Triacylglycerols (TAG).

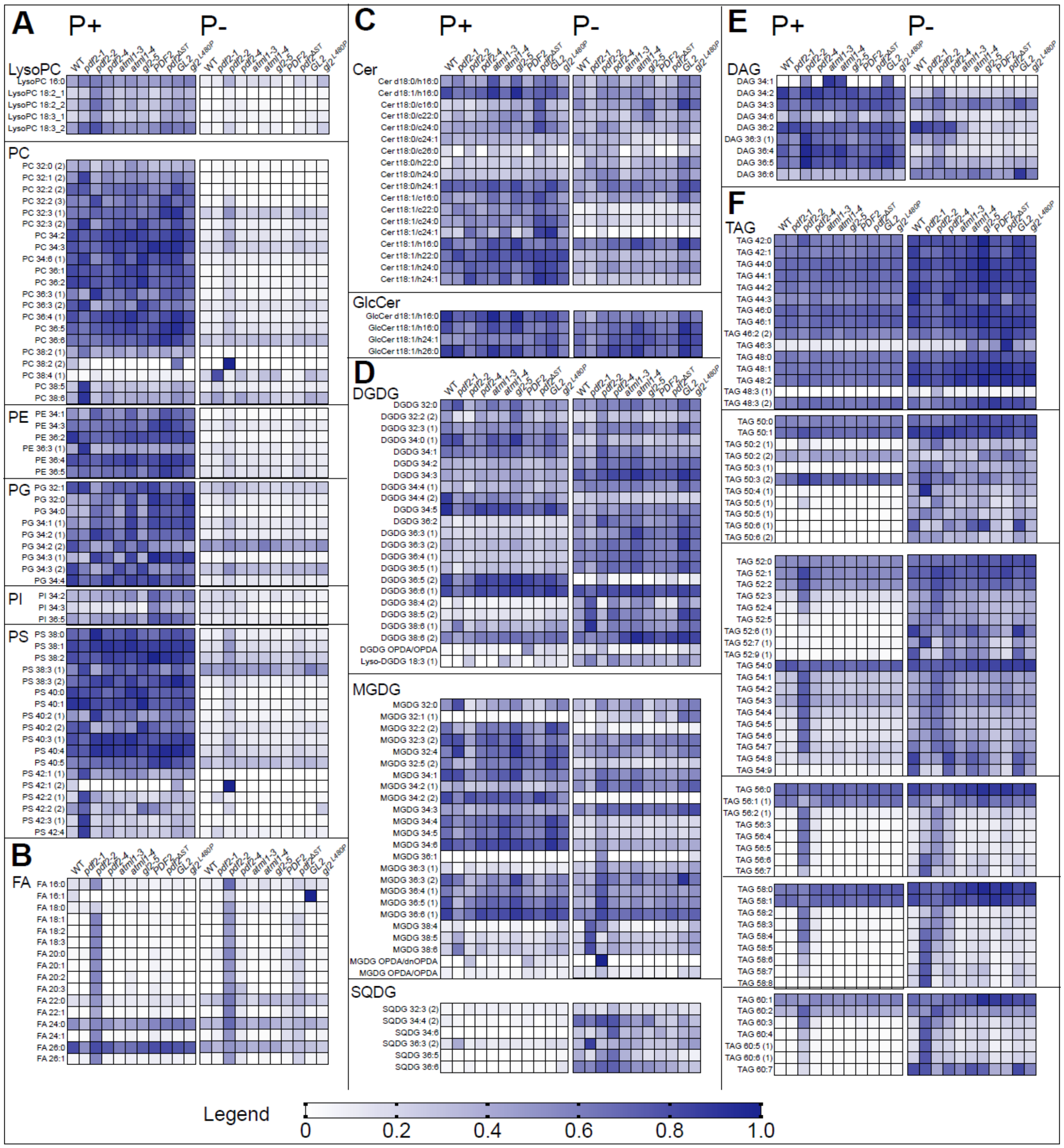


**Supplemental Figure 4.** Heat maps illustrate lipidomic profiles in wild type, *pdf2*, *atml1* and *gl2* mutants under Pi sufficient and limiting conditions. Related to **Figure 4** and **Supplemental Table 4B.**

**(A-F)** Lipidomics data is shown for 14-d-old seedlings whose roots were excised from wild-type (WT), *pdf2-1*, *pdf2-2*, *pdf2-4*, *atml1-3*, *atml1-4* and *gl2-5* genotypes, as well as transgenic lines expressing *EYFP:PDF2* WT, *EYFP:pdf2*$^{\Delta START}$, *EYFP:GL2* WT and *EYFP:gl2*$^{L480P}$ grown in Pi sufficient (P+) (left) and limiting (P-) (right) conditions. Minimum and maximum values were normalized to 0.0 and 1.0, respectively, for visualization purposes. **(A)** Phospholipids (LysoPC, PC, PE, PG, PI, PS), **(B)** Fatty Acids (FA), **(C)** Ceramides (Cer) and glucosylceramides (GlcCer), **(D)** Digalactosyldiacylglycerols (DGDG), monogalactosyldiacylglycerols (MGDG) and sulfoquinovosyl diacylglycerols (SQDG), **(E)** Diacylglycerols (DAG), and **(F)** Triacylglycerols (TAG).

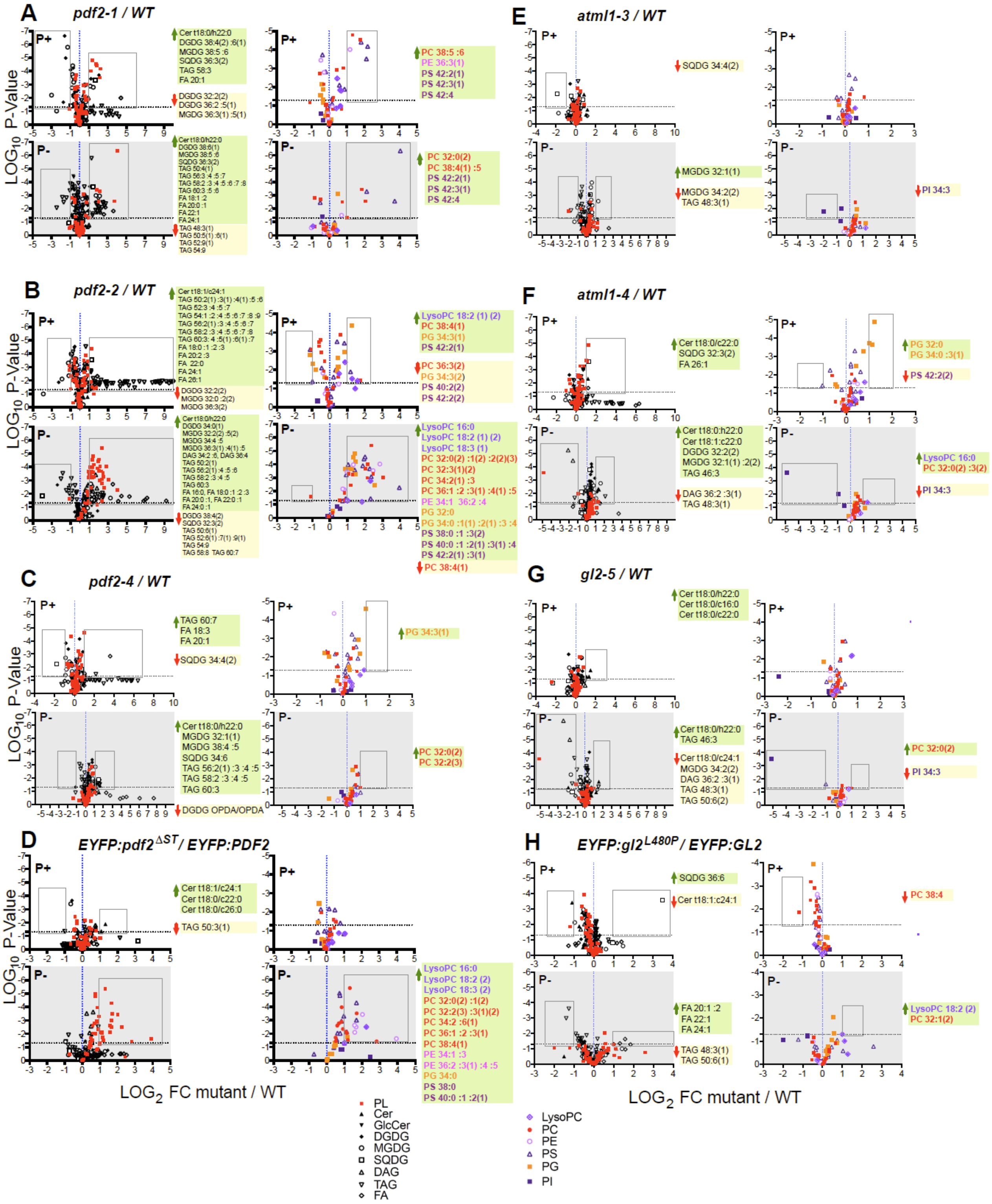
A
pdf2-1 / WT
B
pdf2-2 / WT
C
pdf2-4 / WT
D
EYFP:pdf2ΔST / EYFP:PDF2
E
atml1-3 / WT
F
atml1-4 / WT
G
gl2-5 / WT
H
EYFP:gl2L480P / EYFP:GL2
P+
P-
LOG10 P-Value
LOG2 FC mutant / WT
SQDG 34:4(2)
MGDG 32:1(1)
MGDG 34:2(2)
TAG 48:3(1)
PI 34:3
Cer t18:0/c22:0
SQDG 32:3(2)
FA 26:1
PG 32:0
PG 34:0 :3(1)
PS 42:2(2)
Cer t18:0/h22:0
Cer t18:1:c22:0
DGDG 32:2(2)
MGDG 32:1(1) :2(2)
TAG 46:3
DAG 36:2 :3(1)
LysoPC 16:0
PC 32:0(2) :3(2)
TAG 60:7
FA 18:3
FA 20:1
PG 34:3(1)
PC 32:0(2)
PC 32:2(3)
DGDG OPDA/OPDA
Cer t18:1/c24:1
Cer t18:0/c22:0
Cer t18:0/c26:0
TAG 50:3(1)
Cer t18:0/c16:0
Cer t18:0/c24:1
MGDG 34:2(2)
TAG 50:6(2)
SQDG 36:6
Cer t18:1:c24:1
PC 38:4
FA 20:1 :2
FA 22:1
FA 24:1
TAG 50:6(1)
LysoPC 18:2 (2)
PC 32:1(2)
PL
Cer
GlcCer
DGDG
MGDG
SQDG
DAG
TAG
FA
LysoPC
PC
PE
PS
PG
PI

**Supplemental Figure 5.** Volcano plots reveal lipid changes in HD-Zip IV mutants under Pi sufficient and limiting conditions. Related to **Figure 4, Supplemental Figure 4,** and **Supplemental Table 4C.**

**(A-H)** Volcano plots of general lipid changes (left) versus phospholipid specific changes (right) for mutant versus wild type (WT) are shown. Left: Volcano plots indicate phospholipids (PL) (red), ceramides (Cer), glucosylceramides (GlcCer), digalactosyldiacylglycerols (DGDG), monogalactosyldiacylglycerols (MGDG), sulfoquinovosyl diacylglycerols (SQDG), diacylglyerols (DAG), triacylglycerols (TAG), and fatty acids (FA). Right: Corresponding specific PL changes in phosphatidylcholine (PC), phosphatidylethanolamine (PE), phosphatidylglycerol (PG), phosphoinositol (PI), and phosphatidylserine (PS). Volcano plots depict $\log_2$-fold changes between mutant and WT on the *x* axis versus significance (P values, unpaired *t*-test) on the *y* axis for means of n = 4-5. Horizontal dotted line indicates p = 0.05. Vertical dotted line marks mutant/WT ratio of 1. Lipid species that are upregulated (green arrow) or downregulated (red arrow) relative to WT (FC $\geq$ 2), respectively, are indicated to the right of each graph. Lipid species that are upregulated (green arrow) or downregulated (red arrow) relative to WT (FC $\geq$ 2), respectively, are indicated to the right of each graph. Changes are grouped according to nutrient conditions: Pi sufficient (P+) (top) or limiting (P-) (grey, bottom). Volcano plots depict lipid changes for **(A)** *pdf2-1* versus WT, **(B)** *pdf2-2* versus WT, **(C)** *pdf2-4* versus WT, **(D)** *EYFP:pdf2*$^{\Delta START}$ versus *EYFP:PDF2* WT transgenic lines, **(E)** *atml1-3* versus WT, **(F)** *atml1-4* versus WT, **(G)** *gl2-1* versus WT, and **(H)** *EYFP:gl2*$^{L480P}$ mutant versus *EYFP:GL2* WT transgenic lines.

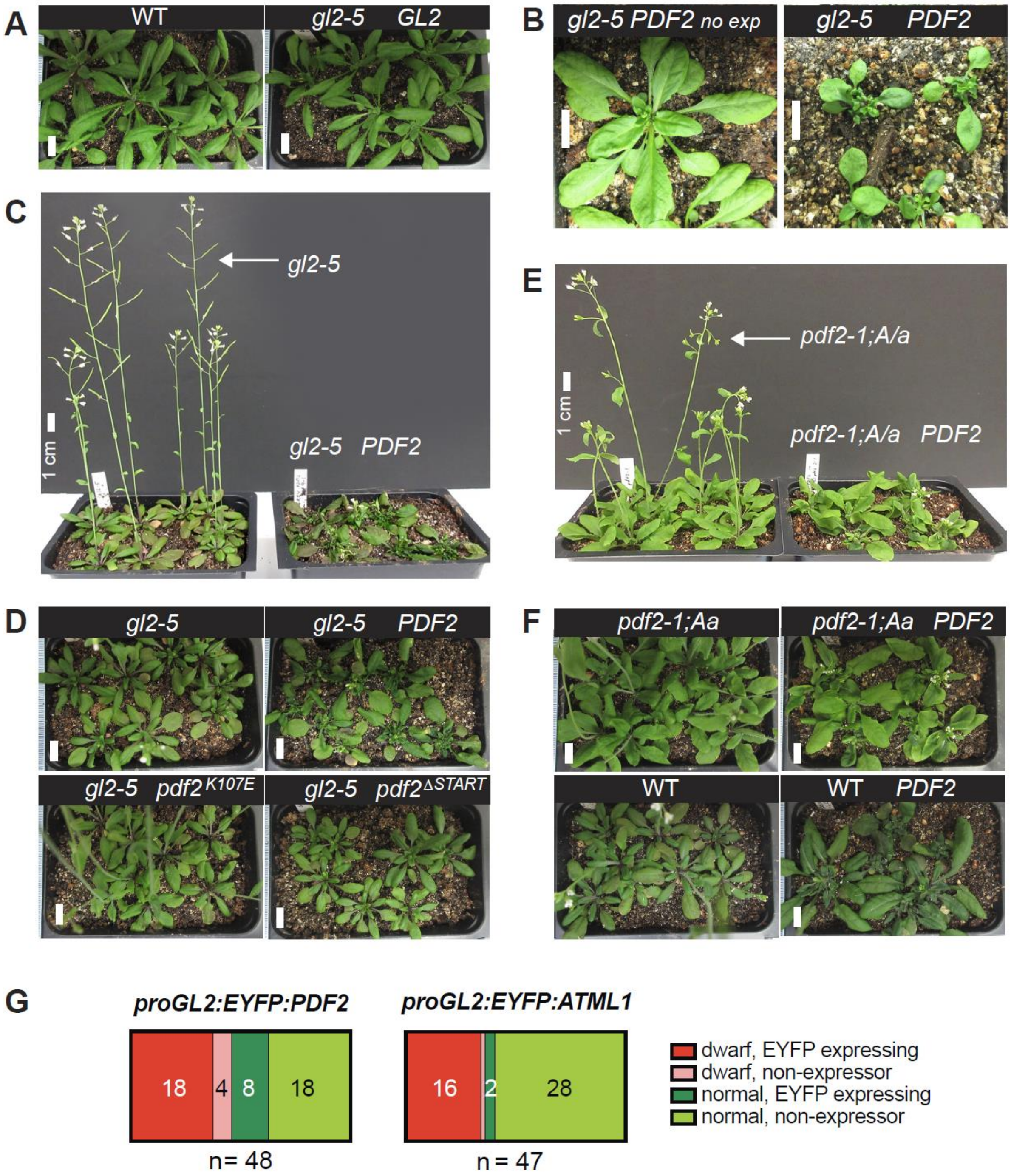


**Supplemental Figure 6.** Plant phenotypes from START-domain dependent expression of PDF2. Related to **Figure 5.**

**(A)** Five-week-old *Arabidopsis* plants of wild type (WT) and transgenic *gl2-5* plants expressing *proGL2:EYFP:GL2* appear similar in morphology. The *proGL2:EYFP:GL2* transgene rescues the *gl2-5* null mutant.

**(B)** Rosettes from transgenic *gl2-5* plants transformed with *proGL2:EYFP:PDF2* having no expression of the transgene (left) versus expression of the transgene (right). Plants expressing the *PDF2* transgene exhibit a dwarf phenotype.

**(C)** Flowering plants of *gl2-5* (left) in comparison to dwarf phenotype of transgenic *gl2-5* plants expressing *proGL2:EYFP:PDF2* (right).

**(D)** Rosette phenotypes of *gl2-5* plants (left) versus *gl2-5* plants expressing *proGL2:EYFP:PDF2* (right) in comparison to *proGL2:EYFP:pdf2*$^{K107E}$ or *proGL2:EYFP:pdf2*$^{\Delta START}$. Wild-type *PDF2*, but not the HD mutant *pdf2*$^{K107E}$ or the START mutant *pdf2*$^{\Delta START}$ exhibit altered morphology.

**(E)** Flowering plants of *pdf2-1;ATML1/atml1-1 (pdf2-1;A/a)* genotype (left) in comparison to dwarf phenotype when this genotype is expressing *proGL2:EYFP:PDF2* (right). Expression of the *proGL2:EYFP:PDF2* in the *pdf2-1;A/a* genetic background results in altered morphology.

**(F)** Rosette phenotypes of *pdf2-1;A/a* plants and WT in comparison to genotypes expressing *proGL2:EYFP:PDF2*. Transgenic lines expressing *proGL2:EYFP:PDF2* in the WT background results in altered morphology. **(A-F)** Size bars = 1 cm.

**(G)** Dwarf phenotype correlates with expression of the *PDF2* or *ATML1* transgene. Plants of genotype *gl2-5* were transformed with *proGL2:EYFP:PDF2* or *proGL2:EYFP:ATML1*. The T1 transformants were classified according to growth phenotype (dwarf or normal) and EYFP expression. Dwarf phenotypes were observed for 44% (21/48) of the *PDF2* and 36% (17/47) of the *ATML1* transformants. Approximately 82% (18/22) and 94% (16/17) of the dwarfed T1 transformants exhibited expression for *PDF2* and *ATML1*, respectively.

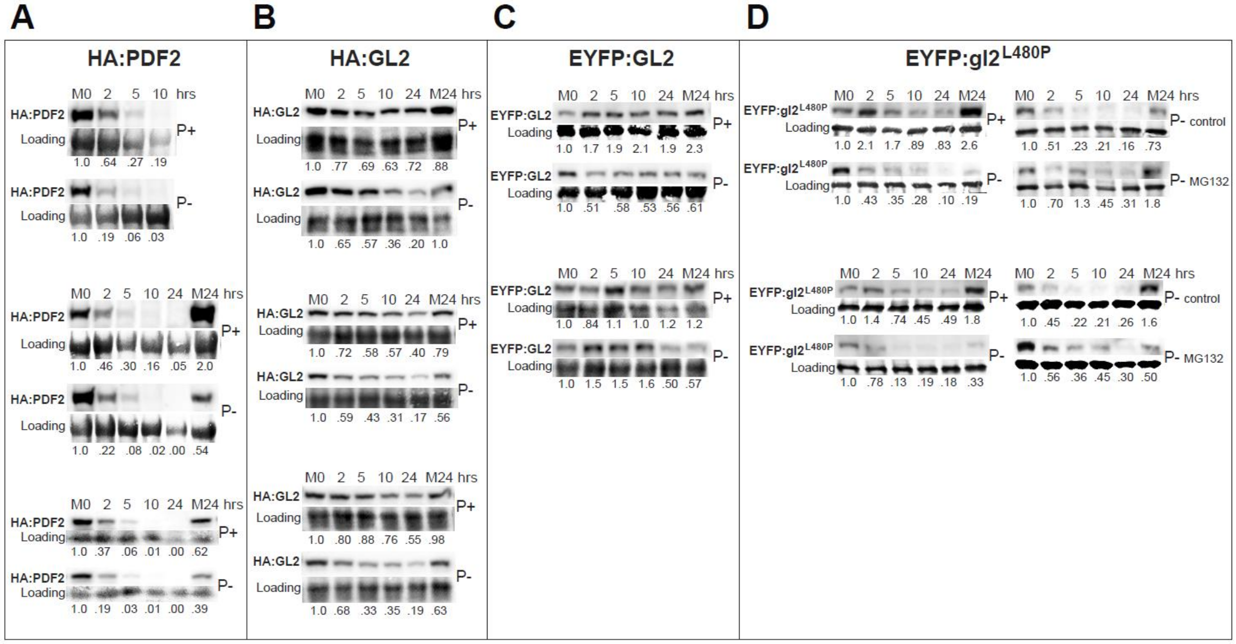


**Supplemental Figure 7.** Protein stability of PDF2 and GL2 is reduced under Pi limitation and START domain mutant exhibits enhanced protein instability. Related to **Figure 6.**

Seedlings were grown either on Pi sufficiency (P+) or limitation (P-) for 5-6 days, followed by cycloheximide treatment (400 µM) for 24 h. Protein levels were monitored by Western blot. Immunodetection with anti-HA **(A and B)** or anti-GFP **(C and D)** antibodies was followed by Coomassie blue staining for loading controls. Top (P+) and bottom (P-) rows are from the same blot. M0 and M24 are DMSO mock treatments at 0 and 24 h, respectively. Representative experiments and protein quantification of Western blots are presented in **Figure 6E and 6F**. Three **(A and B)** or two **(C and D)** additional independent experiments are shown, respectively.

**(A)** Note that the first experiment with HA:PDF2 was conducted for 10 h rather than 24 h. **(A)** HA:PDF2 and **(B)** HA:GL2 exhibited decreased protein levels under Pi limitation.

**(C)** EYFP:GL2 exhibited reduced protein levels that were especially apparent at 24 h.

**(D)** In comparison to wild-type GL2, START domain mutant gl2$^{L480P}$ exhibited a reduction in protein levels that was enhanced under Pi limitation. MG132 treatment (50 µM) resulted in reversal of the observed instability of EYFP:gl2$^{L480P}$.

**Supplemental Table 1.** Lipidomic changes in *atml1-1*;*pdf2-1* vs. wild type. Related to **Supplemental Figures 2 and 3.** Values from **Supplemental Data Set 3** for n = 5 biological replicates were multiple-tested in MetaboAnalyst (Xia et al., 2009) using fold change (FC) $\geq$ 2, adjusted *t*-test P-value $\leq$ 0.05, and FDR correction.

| Lipid | FC up | $LOG_2$ (FC) | P-value |
|---|---|---|---|
| FA 24:1 | 30.556 | 4.9334 | 0.028529 |
| PG 36:4 | 25.2 | 4.6554 | 5.61E-05 |
| Cer t18:0/c24:0 | 22.778 | 4.5096 | 0.012851 |
| TAG 58:5 | 21.924 | 4.4545 | 0.004066 |
| PG 36:5 | 20.833 | 4.3808 | 1.76E-05 |
| TAG 60:5 (1) | 19.556 | 4.2895 | 0.0083778 |
| Cer t18:1/c26:0 | 16 | 4 | 0.039571 |
| TAG 58:6 | 15.645 | 3.9676 | 0.0108 |
| TAG 60:4 | 15.259 | 3.9316 | 0.00076596 |
| Cer t18:0/h24:0 | 15 | 3.9069 | 0.0025998 |
| Cer t18:1/h26:0 | 14 | 3.8074 | 0.028755 |
| FA 18:3 | 13.867 | 3.7935 | 0.00049569 |
| FA 18:1 | 13.426 | 3.7469 | 0.0361 |
| FA 20:1 | 13.02 | 3.7027 | 0.020471 |
| FA 18:2 | 12.705 | 3.6673 | 0.0049023 |
| PE 36:3 1 | 12.256 | 3.6155 | 0.004513 |
| TAG 56:6 | 12.159 | 3.604 | 0.0039214 |
| TAG 56:5 | 12.098 | 3.5967 | 0.0025934 |
| Cer t18:1/c16:0 | 12 | 3.585 | 0.048221 |
| Cer t18:1/h24:1 | 11.709 | 3.5496 | 0.00068263 |
| TAG 58:4 | 11.639 | 3.5409 | 0.00032303 |
| Cer d18:0/h16:0 | 11 | 3.4594 | 0.040911 |
| FA 22:0 | 10.818 | 3.4354 | 0.0018722 |
| Cer t18:0/c24:1 | 10 | 3.3219 | 0.0098589 |
| FA 20:0 | 9.6383 | 3.2688 | 0.0040639 |
| TAG 56:7 | 9.5567 | 3.2565 | 0.0032396 |
| TAG 58:7 | 9.4 | 3.2327 | 0.0098582 |
| LysoPC 18:3 (1) | 9 | 3.1699 | 0.022501 |
| Cer t18:1/h26:1 | 8.3333 | 3.0589 | 0.012981 |
| Cer t18:0/h22:0 | 8 | 3 | 0.0098582 |
| PC 32:3 (3) | 8 | 3 | 0.028841 |
| TAG 58:3 | 7.96 | 2.9928 | 0.0010919 |
| PC 32:3 (1) | 7.9091 | 2.9835 | 0.00319 |
| PG 36:6 | 7.82 | 2.9672 | 9.72E-05 |
| TAG 56:4 | 7.7278 | 2.9501 | 0.00097797 |
| TAG 52:9 (1) | 7.2 | 2.848 | 0.0108 |
| TAG 60:3 | 6.8049 | 2.7666 | 0.0025883 |
| Cer t18:1/h24:0 | 6.5732 | 2.7166 | 0.0028019 |
| FA 16:0 | 6.543 | 2.71 | 0.0064766 |
| TAG 60:7 | 6.5 | 2.7004 | 0.0025235 |
| Cer t18:1/c24:1 | 6.4 | 2.6781 | 0.0039511 |
| Cer t18:1/h22:0 | 6.2619 | 2.6466 | 0.0032594 |
| PG 36:3 | 6.1429 | 2.6189 | 0.0028019 |
| TAG 54:9 | 6.0411 | 2.5948 | 0.0048124 |
| FA 18:0 | 5.8955 | 2.5596 | 0.012241 |
| TAG 54:8 | 5.6654 | 2.5022 | 0.004513 |
| TAG 56:3 | 5.3916 | 2.4307 | 0.000923 |
| PC 32:1 (1) | 5.1818 | 2.3735 | 0.028755 |
| TAG 54:7 | 5.1653 | 2.3689 | 0.0098873 |
| PG 34:3 (1) | 4.8588 | 2.2806 | 2.33E-07 |
| PG 34:0 | 4.3696 | 2.1275 | 0.0010023 |
| TAG 56:8 | 4.25 | 2.0875 | 0.00031619 |
| TAG 52:6 (1) | 3.649 | 1.8675 | 0.0026739 |
| PG 32:0 | 3.5747 | 1.8378 | 2.94E-05 |
| PE 36:6 | 3.4405 | 1.7826 | 0.0066656 |
| TAG 56:2 (1) | 2.9892 | 1.5798 | 0.0053121 |
| TAG 52:5 | 2.8571 | 1.5146 | 0.0060647 |
| PS 40:5 | 2.6605 | 1.4117 | 0.00068263 |
| TAG 54:6 | 2.6374 | 1.3991 | 0.0053121 |
| PG 34:2 (1) | 2.6099 | 1.384 | 0.00055821 |
| PG 34:1 (1) | 2.3572 | 1.2371 | 0.0010768 |
| PE 36:5 | 2.3527 | 1.2343 | 0.033137 |
| PI 36:6 | 2.34 | 1.2265 | 0.0014895 |
| DAG 36:6 | 2.2222 | 1.152 | 0.00319 |
| PC 36:6 | 2.0597 | 1.0425 | 0.0091884 |

| Lipid | FC dn | $LOG_2$ (FC) | P-value |
|---|---|---|---|
| MGDG 34:6 | 0.43495 | -1.2011 | 0.0010023 |
| DGDG 36:6 | 0.41357 | -1.2738 | 0.000923 |
| DGDG 38:6 (2) | 0.41264 | -1.277 | 0.0025934 |
| SQDG 36:4 (1) | 0.41194 | -1.2795 | 0.0003254 |
| SQDG 34:4 (2) | 0.40541 | -1.3026 | 0.03452 |
| DGDG 34:2 | 0.39102 | -1.3547 | 0.0012811 |
| PC 32:1 (2) | 0.37838 | -1.4021 | 0.0028019 |
| MGDG 36:6 (1) | 0.37568 | -1.4124 | 0.0001845 |
| MGDG 32:4 | 0.36585 | -1.4507 | 0.0008783 |
| DGDG 34:3 | 0.36202 | -1.4658 | 0.0013534 |
| MGDG 34:4 | 0.36074 | -1.471 | 0.0012686 |
| MGDG 34:3 | 0.35866 | -1.4793 | 0.0006826 |
| GlcCer d18:1/h16:0 | 0.34936 | -1.5172 | 0.0005582 |
| SQDG 36:3 (2) | 0.3439 | -1.5399 | 0.0003859 |
| DGDG 32:0 | 0.34228 | -1.5467 | 0.000923 |
| GlcCer t18:1/h16:0 | 0.33546 | -1.5758 | 0.0003230 |
| GlcCer t18:1/h26:0 | 0.31486 | -1.6672 | 0.0004178 |
| DGDG 36:3 (1) | 0.29091 | -1.7814 | 0.021145 |
| Cer d18:1/h16:0 | 0.28896 | -1.7911 | 0.0007779 |
| MGDG 34:2 (1) | 0.25748 | -1.9575 | 0.0001867 |
| Cer t18:1/h16:0 | 0.25704 | -1.9599 | 0.0001431 |
| DGDG 34:0 (1) | 0.25168 | -1.9903 | 0.0015093 |
| DGDG 38:6 (1) | 0.24563 | -2.0255 | 0.0003254 |
| MGDG 38:6 | 0.2351 | -2.0886 | 0.0001431 |
| DGDG 36:5 (2) | 0.22857 | -2.1293 | 0.021517 |
| MGDG 32:3 (2) | 0.21622 | -2.2095 | 0.0006600 |
| DGDG 34:5 | 0.21474 | -2.2193 | 0.0001867 |
| SQDG 36:5 | 0.21278 | -2.2326 | 5.70E-06 |
| PC 36:1 | 0.20556 | -2.2824 | 0.004513 |
| SQDG 36:3 (1) | 0.20492 | -2.2869 | 0.0003230 |
| PG 36:7 | 0.19012 | -2.395 | 0.003371 |
| DGDG 36:5 (1) | 0.18703 | -2.4187 | 0.0001801 |
| DGDG 34:4 (2) | 0.18462 | -2.4374 | 0.002051 |
| SQDG 34:2 | 0.18242 | -2.4547 | 5.70E-06 |
| MGDG 34:5 | 0.18102 | -2.4658 | 0.0005582 |
| DGDG 32:3 (1) | 0.17722 | -2.4964 | 0.0004735 |
| SQDG 34:6 | 0.17586 | -2.5075 | 0.0025998 |
| PG 34:3 (2) | 0.17266 | -2.534 | 0.0028019 |
| MGDG 32:0 | 0.16667 | -2.585 | 0.01433 |
| DGDG 36:3 (2) | 0.15 | -2.737 | 0.0006826 |
| MGDG 36:5 (1) | 0.14057 | -2.8307 | 0.0003162 |
| GlcCer t18:1/h20:0 | 0.13725 | -2.8651 | 0.0034767 |
| MGDG 32:2 (2) | 0.13514 | -2.8875 | 0.0038123 |
| MGDG 32:5 (2) | 0.12069 | -3.0506 | 0.0005582 |
| MGDG 36:3 (2) | 0.12037 | -3.0544 | 0.0067979 |
| SQDG 34:5 | 0.1125 | -3.152 | 0.0024905 |
| DGDG 38:5 (2) | 0.11124 | -3.1683 | 0.0001801 |
| MGDG 36:3 (1) | 0.090909 | -3.4594 | 0.040911 |
| MGDG 38:5 | 0.074286 | -3.7508 | 0.0002733 |
| DGDG 36:2 | 0.045763 | -4.4497 | 0.0028019 |

**Supplemental Table 2.** Lipidomic changes in *pdf2-1* vs. wild type under Pi limitation. Related to **Figure 4.** Values from **Supplemental Data Set 4** for n = 5 biological replicates each for *pdf2-1* and wild type were multiple-tested for significance in MetaboAnalyst (Xia et al., 2009) using fold change (FC) $\geq$ 2, adjusted *t*-test P-value $\leq$ 0.05, and FDR correction.

| Lipid | FC | $LOG_2$ (FC) | P-value |
|---|---|---|---|
| MGDG 38:4 | 46.096 | 5.5266 | 0.013781 |
| PS 42:3 (1) | 26.307 | 4.7174 | 5.13E-05 |
| PS 42:2 (1) | 13.197 | 3.7221 | 0.013283 |
| FA 18:1 | 12.352 | 3.6266 | 0.035422 |
| PC 32:1 (2) | 10.463 | 3.3872 | 0.034313 |
| FA 24:1 | 10.226 | 3.3541 | 0.016768 |
| FA 22:1 | 8.329 | 3.0581 | 0.020773 |
| TAG 58:6 | 6.9645 | 2.8 | 0.015066 |
| TAG 58:5 | 6.4617 | 2.6919 | 0.018918 |
| FA 20:1 | 6.0202 | 2.5898 | 0.013781 |
| TAG 60:4 | 5.6696 | 2.5033 | 0.01408 |
| TAG 60:5 (1) | 5.577 | 2.4795 | 0.013781 |
| TAG 58:4 | 5.3317 | 2.4146 | 0.022647 |
| TAG 58:3 | 4.4815 | 2.164 | 0.035911 |
| MGDG 38:5 | 4.4417 | 2.1511 | 0.0080159 |
| PS 42:4 | 4.368 | 2.127 | 0.0055989 |
| TAG 60:3 | 4.2824 | 2.0984 | 0.022647 |
| TAG 56:6 | 4.1907 | 2.0672 | 0.027236 |
| FA 20:0 | 4.1571 | 2.0556 | 0.018918 |
| PC 38:4 (1) | 4.1185 | 2.0421 | 0.013283 |
| TAG 58:8 | 4.0804 | 2.0287 | 0.010222 |
| TAG 56:5 | 3.7365 | 1.9017 | 0.035276 |
| TAG 56:7 | 3.6789 | 1.8793 | 0.016942 |
| TAG 60:6 (1) | 3.6097 | 1.8519 | 0.01408 |
| PC 38:5 | 3.5897 | 1.8439 | 0.0046714 |
| TAG 58:7 | 3.4288 | 1.7777 | 0.015066 |
| MGDG 34:2 (2) | 3.4055 | 1.7678 | 0.013283 |
| MGDG 38:6 | 3.3777 | 1.756 | 0.00788 |
| TAG 56:4 | 3.2923 | 1.7191 | 0.031992 |
| SQDG 36:3 (2) | 2.7372 | 1.4527 | 0.0041771 |
| TAG 50:4 (1) | 2.426 | 1.2786 | 0.00017736 |
| DGDG 38:6 (1) | 2.1173 | 1.0823 | 0.0041771 |
| TAG 50:5 (1) | 0.49025 | -1.0284 | 0.0050232 |
| Cer d18:0/h16:0 | 0.46637 | -1.1004 | 0.031992 |
| TAG 52:9 (1) | 0.37982 | -1.3966 | 0.0094223 |
| TAG 50:6 (1) | 0.30819 | -1.6981 | 0.013781 |
| TAG 54:9 | 0.27969 | -1.8381 | 0.0094223 |
| SQDG 34:6 | 0.22541 | -2.1494 | 0.0072859 |
| TAG 48:3 (1) | 0.18455 | -2.4379 | 0.023821 |
| MGDG 32:1 (1) | 0.13104 | -2.9319 | 0.0072859 |

**Supplemental Table 3.** Lipidomic changes in *pdf2-2* vs. wild type under Pi limitation. Related to **Fig. 4.** Values from **Supplemental Data Set 4** for n = 5 biological replicates each for *pdf2-2* and wild type (WT) were multiple-tested for significance in MetaboAnalyst (Xia et al., 2009) using fold change (FC) $\geq$ 2, adjusted *t*-test P-value $\leq$ 0.05, and FDR correction.

| Lipid | FC | $LOG_2$ (FC) | P-value |
|---|---|---|---|
| PC 32:1 (2) | 101.06 | 6.6591 | 0.035619 |
| FA 24:1 | 86.136 | 6.4285 | 0.044978 |
| FA 26:1 | 63.207 | 5.982 | 0.021511 |
| PS 42:3 (1) | 24.86 | 4.6358 | 0.022407 |
| PS 40:0 | 24.076 | 4.5895 | 0.011039 |
| FA 20:0 | 21.425 | 4.4212 | 0.044862 |
| PC 32:2 (3) | 19.82 | 4.3089 | 0.0093426 |
| MGDG 32:2 (2) | 19.207 | 4.2636 | 0.048343 |
| LysoPC 18:2 (1) | 14.605 | 3.8684 | 0.042562 |
| FA 16:0 | 11.406 | 3.5118 | 0.011863 |
| PC 36:1 | 8.1446 | 3.0258 | 0.0070196 |
| PC 32:0 (2) | 7.3406 | 2.8759 | 0.014694 |
| PE 34:1 | 7.1753 | 2.843 | 0.0024257 |
| PG 34:0 | 5.9078 | 2.5626 | 0.017529 |
| TAG 56:4 | 5.7133 | 2.5143 | 0.044978 |
| TAG 56:3 | 5.7089 | 2.5132 | 0.041479 |
| PC 32:3 (2) | 5.5473 | 2.4718 | 0.0051214 |
| PE 36:2 | 5.5176 | 2.464 | 0.0036406 |
| TAG 58:2 | 5.2898 | 2.4032 | 0.032056 |
| PC 36:3 (1) | 5.1346 | 2.3603 | 0.00087379 |
| PS 40:1 | 5.1289 | 2.3587 | 0.0065128 |
| TAG 58:3 | 5.0829 | 2.3457 | 0.041327 |
| LysoPC 16:0 | 4.9686 | 2.3128 | 0.011594 |
| PS 38:0 | 4.8303 | 2.2721 | 0.0032786 |
| PC 34:2 | 4.4435 | 2.1517 | 0.0024257 |
| PC 36:2 | 4.3875 | 2.1334 | 0.007537 |
| PC 36:4 (1) | 4.075 | 2.0268 | 0.0032786 |
| PS 40:2 (1) | 4.0193 | 2.0069 | 0.0015725 |
| PS 38:1 | 3.8608 | 1.9489 | 0.0017704 |
| TAG 56:2 (1) | 3.7379 | 1.9022 | 0.035619 |
| PS 40:3 (1) | 3.6796 | 1.8795 | 0.0026047 |
| DAG 34:6 | 3.5084 | 1.8108 | 0.018283 |
| DAG 36:4 | 3.2076 | 1.6815 | 0.0041482 |
| TAG 60:3 | 3.0982 | 1.6315 | 0.041141 |
| SQDG 32:3 (2) | 2.9393 | 1.5555 | 0.042676 |
| FA 22:0 | 2.8484 | 1.5102 | 0.04963 |
| PG 34:2 (1) | 2.787 | 1.4787 | 0.0051214 |
| PG 34:3 (1) | 2.7562 | 1.4627 | 0.0024257 |
| PG 34:4 | 2.6884 | 1.4268 | 0.0015725 |
| PS 38:3 (2) | 2.6748 | 1.4195 | 0.0034835 |
| DAG 34:2 | 2.6154 | 1.3871 | 0.026837 |
| PE 36:4 | 2.5293 | 1.3387 | 0.0024257 |
| PG 32:0 | 2.5042 | 1.3244 | 0.0015725 |
| PC 36:5 | 2.4557 | 1.2962 | 0.0052882 |
| MGDG 34:5 | 2.4401 | 1.2869 | 0.014694 |
| PG 34:1 (1) | 2.3825 | 1.2525 | 0.0032786 |
| DGDG 34:0 (1) | 2.3055 | 1.2051 | 0.026893 |
| PC 32:3 (1) | 2.2953 | 1.1987 | 0.017529 |
| MGDG 34:4 | 2.2771 | 1.1872 | 0.014694 |
| MGDG 36:5 (1) | 2.2391 | 1.1629 | 0.011668 |
| MGDG 36:4 (1) | 2.1529 | 1.1063 | 0.026893 |
| PC 34:3 | 2.1334 | 1.0932 | 0.0050838 |
| FA 24:0 | 2.0865 | 1.0611 | 0.045749 |
| PS 40:4 | 2.0639 | 1.0454 | 0.0024257 |
| PG 34:3 (2) | 2.0041 | 1.003 | 0.0032786 |
| TAG 60:7 | 0.49992 | -1.0002 | 0.014605 |
| TAG 58:8 | 0.45946 | -1.122 | 0.034248 |
| TAG 52:7 (1) | 0.45379 | -1.1399 | 0.018283 |
| TAG 50:6 (1) | 0.36744 | -1.4444 | 0.024956 |
| TAG 52:9 (1) | 0.28401 | -1.816 | 0.0036406 |
| TAG 54:9 | 0.24826 | -2.0101 | 0.006361 |
| DGDG 38:4 (2) | 0.1901 | -2.3952 | 0.013126 |

**Supplemental Table 4.** Lipidomic changes in *EYFP:pdf2*$^{\Delta START}$ vs. *EYFP:PDF2* under Pi limitation. Related to **Figure 4.** Values from **Supplemental Data Set 4** for n = 4 biological replicates each for *pdf2*$^{\Delta START}$ and *PDF2* were multiple-tested for fold-change differences in MetaboAnalyst (Xia et al., 2009) using fold change (FC) $\geq$ 2, adjusted *t*-test P-value $\leq$ 0.05, and FDR correction.

| **Lipid** | **FC** | **LOG$_2$ (FC)** | **P-value** |
|---|---|---|---|
| PE 36:2 | 15.221 | 3.9279 | 0.040654 |
| LysoPC 18:2 (2) | 9.4602 | 3.2419 | 0.017649 |
| PC 32:2 (3) | 7.9487 | 2.9907 | 0.028085 |
| LysoPC 16:0 | 4.7111 | 2.2361 | 0.0092167 |
| PE 36:3 (1) | 4.0439 | 2.0157 | 0.0044461 |
| PC 36:3 (1) | 3.2245 | 1.6891 | 0.0029955 |
| PS 40:2 (1) | 3.2036 | 1.6797 | 0.0019275 |
| PS 38:0 | 3.1974 | 1.6769 | 0.0012466 |
| PE 34:3 | 3.1488 | 1.6548 | 0.0092167 |
| PE 36:4 | 2.9868 | 1.5786 | 0.017308 |
| PE 36:5 | 2.9715 | 1.5712 | 0.0066889 |
| PC 32:3 (1) | 2.4704 | 1.3047 | 0.00057037 |
| PC 36:2 | 2.1914 | 1.1319 | 0.028085 |
| PS 40:1 | 2.1904 | 1.1312 | 0.033294 |
| PC 34:2 | 2.0937 | 1.0661 | 0.02873 |

**Supplemental Table 5.** Oligonucleotides used in this study.

| | |
|---|---|
| I. Primers for cloning into pKCTAP, pKNGSTAP, and pET-His6-MBP-TEV-LIC. Gene-specific sequences are | |
| PDF2 TAP N' F | GGG GAC AAG TTT GTA CAA AAA AGC AGG CGT G **ATG TAC CAT CCA AAC ATG TTT GAG** |
| PDF2 TAP N' R | GGG GAC CAC TTT GTA CAA GAA AGC TGG GT **CTA CGC TCC TCC TCC AAC ATC** |
| PDF2 TAP C' F | GGG GAC AAG TTT GTA CAA AAA AGC AGG CGC CACC **ATG TAC CAT CCA AAC ATG TTT** |
| PDF2 TAP C' R | GGG GAC CAC TTT GTA CAA GAA AGC TGG GTG **CGC TCC TCC TCC AAC ATC AC** |
| PDF2_MBP_LIC_F | TACTTCCAATCCAATGCA **GGA ACA GGG GAC ATT TTG AG** |
| PDF2 MBP LIC R | TTATCCACTTCCAATGTTATTA **CAT GGA GCT AGC AAG CCG** |
| PDF2 ΔSTART F | CGG CTT GCT AGC TCC ATG GC |
| PDF2 ΔSTART R | ATC AGT CTC AGA AGG AAT CGA AAC |
| II. Primers for genotyping insertion mutants. Mutant and wild-type alleles were detected in 3-primer PCRs. | |
| atml1-3 F (oAR272) | CAGGCAGAAGAAAATCGAGAT (Meyer et al., 2017) |
| atml1-3 R (oAR273) | GAAACCAGTGTGGCTATTGTT (Meyer et al., 2017) |
| LBb1 (T-DNA border for *atml1-3*) | GCGTGGACCGCTTGCTGCAACT |
| atml1-4 F (oAR274) | TACGTAGGGAAGCCTTTAATG |
| atml1-4 R (oAR275) | CATACCGGAAAGATCACAAG |
| JMLB2 (T-DNA border for *atml1-4*) | TTGGGTGATGGTTCACGTAGTGGG |
| atml1-1 F (ML1-F1) | TGGGATATACAGGCAGAAGA (Ogawa et al., 2015) |
| atml1-1 R (ML1-3N) | GTTTTGGAGCTACAGGGATCCAGA (Ogawa et al., 2015) |
| pdf2-1 F (PDF2 F1) | GATCAGTGCCTTGAAGGAAA (Ogawa et al., 2015) |
| pdf2-1 R (PDF2 R2) | CTGTTGTCGACATTGTTGTC (Ogawa et al., 2015) |
| JL202 (T-DNA border for *atml1-1* and | CATTTTATAATAACGCTGCGGACATCTAC (Ogawa et al., 2015) |
| pdf2-2 F | CTCGTTCGCTTGATCTTGAAG |
| pdf2-2 R | AGATCACCAGGAATGTTGCTG |
| LBb1.3 (T-DNA border for *pdf2-2*) | ATTTTGCCGATTTCGGAAC |
| pdf2-4 F | TGCCTCATTTTTAACAAATATTATCG |
| pdf2-4 R | TCTGGTTCTCATGCCTCTCAC |
| LB1 SAIL (T-DNA border for *pdf2-4*) | GCCTTTTCAGAAATGGATAAATAGCCTTGCTTCC |
| gl2-5 F (GL2 F 112) | ATGTCAATGGCCGTCGACATGTC (Wang et al., 2007) |
| gl2-5 R (GL2 R 1462) | TCTCGCAGCTTCTCTAGTTCCG (Wang et al., 2007) |
| En8130 (T-DNA border for *gl2-5*) | GAGCGTCGGTCCCCACACTTCTATAC (Baumann et al., 1998) |
| III. Primers to generate K107E and L467P missense mutations and START deletion in *PDF2*, and L480P missense mutation in *GL2*. Codons for glutamic acid (E) and proline (P) are indicated in bold. | |
| pdf2 K107E F | CT CTT CAA GTT **GAG** TTT TGG TTC C |
| pdf2 K107E R | G CTC TAA ATT GAG ATC ACG GC |
| pdf2 L467P F | CGT TGG GTG GCT ACA **CCA** GAA CGA CAA TGC GAG |
| pdf2 L467P R | CTC GCA TTG TCG TTC **TGG** TGT AGC CAC CCA ACG |
| pdf2ΔSTART F | GCT AGC TCC ATG GCC AGC |
| pdf2ΔSTART R | AGA AGG AAT CGA AAC TGA CCT C |
| gl2 L480P F | G GTC GCC ACC **CCT** CAG CTC CAT TGC G |
| gl2 L480P R | C GCA ATG GAG CTG **AGG** GGT GGC GAC C |
| IV. Oligonucleotides used in electrophoretic mobility shift assay (EMSA). | |
| PDF2_L1_EMSA_F_Cy3 | /5Cy3/AAA **GAATATTC GAATATTC GAATATTC** CAA |
| PDF2_L1_EMSA_F | AAA **GAATATTC GAATATTC GAATATTC** CAA |
| PDF2_L1_EMSA_R | TTG **GAATATTC GAATATTC GAATATTC** TTT |
| V. Primers used in qRT-PCR experiments | |
| GL2_PCR_F | TGGATTGCCACTGAGTTGCCTCTG (amplifies 106) (Ishida et al., 2007) |
| GL2_PCR_R | TGGGGCTGATCGTTGGGTTACCA (Ishida et al., 2007) |
| GDPD1_RTPCR(128)_F | TCTTGTCAGGAAGTTGCAGACC (amplifies 128 bp) |
| GDPD1_RTPCR(128)_R | GGAGGCCTCCTTCTAAACATAC |
| GDPD2_RTPCR(124)_F | TCAGACCGTCTATGAACGAGAG (amplifies 124 bp) |
| GDPD2_RTPCR(124)_R | AAGCTTAGCTGCATCTGGTTGG |
| GDPD3_RTPCR(110)_F | GCGGCTCGACTCATCAGG (amplifies 110 bp) |
| GDPD3_RTPCR(110)_R | ATGGCCTCGTCTAATGAGTTCC |
| NPC2_RTPCR(133)_F | CGGCTTCTGATGATCATCCATC (amplifies 133 bp) |
| NPC2_RTPCR(133)_R | CCACCATGCTCGTCGTAAGTG |
| NPC4_RTPCR(87)_F | GGGTTCCGACTTTCTTCATCTC (amplifies 87 bp) |
| NPC4_RTPCR(87)_R | CATATTGTGACCTTGGGTACGG |
| NPC6_RTPCR(100)_F | GGTTAGGTGTTCGTGTTCCTAC (amplifies 100 bp) |
| NPC6_RTPCR(100)_R | TCGTACTCTGAGCTCTCTGTTG |
| PDF2_RTPCR(185)_F | GGGTTTCACTGTGTCTCC (amplifies 185 bp) |
| PDF2_RTPCR_R | CAAACATGTTTGGATGGTAC (Kamata et al., 2013) |
| SR54_EYFP_F | GTCCTGCTGGAGTTCGTG (amplifies 125 bp in transgenic lines) |
| PDF2_RTPCR(185)_R | GCTCTCAAACATGTTGGATGG |
| PLDe_RTPCR(90)_F | GTCCACTCCAAGCTCATGATAG (amplifies 90 bp) |
| PLDe_RTPCR(90)_R | GTCCCGACAACCATCCATT |
| PLDz1_RTPCR(137)_F | TGGATGGCAACCGCAAAGACAA (amplifies 137 bp) (Li et al., 2006) |
| PLDz1_RTPCR(137)_R | ATCGTTGTGTGTCCCAGCTTCT (Li et al., 2006) |

| | |
|---|---|
| PLDz2_RTPCR(132)_F | ACGATGTCCACTGCGCTCTATG (amplifies 132 bp) (Su et al., 2018) |
| PLDz2_RTPCR(132)_R | GGTGGTGAGGCATCAACAATGG (Su et al., 2018) |
| ACT7_PCR(94)_F | TCGCACATGTACTCGTTTCGCTTTC (amplifies 94 bp) (Khosla et al., 2014) |
| ACT7_PCR(94)_R | TCGAGAAGCAGCGAGAGAGAAAGATAGA (Khosla et al., 2014) |
| VI. Primers to generate *proGL2:EYFP:PDF2* and *proGL2:EYFP:ATML1* constructs using DNA assembly | |
| SR54 EYFP:PDF2 F | CAAGCTTCGAATTCTGCAGTCGACATGTACCATCCAAACATGTTTGAG |
| SR54 PDF2 R | ACCGCGGTGGAGCTCGCTACGCTCCTCCTCCAAC |
| SR54 EYFP:ATML1 F | TCAAGCTTCGAATTCTGCAGTCGACATGTATCATCCAAACATGTTCTC |
| SR54 ATML1 R | CACCGCGGTGGAGCTCGTTAGGCTCCGTCGCAGGCC |

### Supplemental Data Sets in xls spreadsheet (separate document)

Supplemental Data Sets 1A and 1B (xls)
Supplemental Data Sets 2A and 2B (xls)
Supplemental Data Sets 3A and 3B (xls)
Supplemental Data Sets 4A, 4B, and 4C (xls)
Supplemental Data Sets 5A, 5B and 5C (xls)

**Supplemental Data Set 1A.** TAP lipidomics data for the PDF2 TF from soluble fraction. Related to **Figure 1**. Full-length PDF2 TF was compared to pdf2ΔSTART and the empty vector (EV) control. Soluble fraction served as the input material. Data are expressed as log2 of the measured intensity. Given are peak name, annotation, m/z, retention time (RT), and ionization mode.

**Supplemental Data Set 1B.** TAP lipidomics data for the PDF2 TF from soluble fraction. Related to **Figure 1**. Full-length PDF2 TF was compared to pdf2ΔSTART and the empty vector (EV) control. Data are expressed as log2 of the ratio calculated between eluate and input. T-test and FC were calculated only if data were available for three or more replicates. If a molecule was found in none or one of the samples of one line but 5 or 6 replicates of the other line, the P-value and FC were assigned as 0.05 and 2, respectively. Given are peak name, annotation, m/z, retention time (RT), and ionization mode.

**Supplemental Data Set 2A.** List of putative transcriptional target genes from DAP-seq data for PDF2. Related to **Figure 3**. Arabidopsis ID for PDF2 (AT4G04890) is listed alongside IDs for putative target genes.

**Supplemental Data Set 2B.** GO enrichment for DAP-seq targets of PDF2. Related to **Figure 3**. The list of putative target genes was queried for Gene Ontology (GO) enrichment of 'biological process' using the PANTHER Overrepresentation Test. 'Phospholipid catabolic process' and 'cellular response to phosphate starvation' represent the top two categories, with fold enrichment of ~9.8 and ~6.3, respectively.

**Supplemental Data Set 3A.** Comprehensive lipidomic data from wild-type and *pdf2*, *atml1;pdf2*, *atml1*, and *gl2* mutants. Related to **Suppplemental Figure 2**,

**Supplemental Figure 3** and **Supplemental Table 1**. Given are cluster name, m/z, retention time (RT), compound ID, lipid classification and intensities for each sample. Averages and standard deviations were calculated from 5 biological replicates. Intensities were summed for molecules from each lipid classification. T-test and FC were calculated only if data were available for three or more replicates. FC values and corresponding log2 values are given for each of the mutants over WT. P-values were calculated from unpaired t-tests. If a molecule was found in one or none of the samples in WT but in 5 replicates in the mutant, the FC and P-value were assigned as 2 and 0.05, respectively. Color scheme: FC>2, light red; FC>4, red; FC<2, light blue; FC<4, blue (P-value <0.05).

**Supplemental Data Set 3B**. Maximum normalized comprehensive lipidomic data from wild-type and *pdf2*, *atml1;pdf2*, *atml1*, and *gl2* mutants. Related to **Supplemental Figure 3**. Given are cluster name, m/z, retention time (RT), compound ID, lipid classification and intensities for each sample. Averages and standard deviations were calculated from n = 5 biological replicates. Minimum and maximum values were normalized to 0.0 and 1.0, respectively, for visualization purposes in heat maps.

**Supplemental Data Set 4A**. Lipidomic data from wild-type, *pdf2*, *atml1*, and *gl2* mutants, as well as *EYFP:PDF2*, *EYFP:pdf2*$^{\Delta START}$, *EYFP:GL2*, and *EYFP:gl2*$^{L480P}$ transgenic lines in Pi sufficient (P+) and Pi limiting (P-) media. Related to **Figure 4** and **Supplemental Tables 2-4**. Given are cluster name, m/z, retention time (RT), compound ID, lipid classification, adduct, and intensities for each sample. Averages and standard deviations were calculated from n = 4 or 5 biological replicates. Intensities were summed for molecules from each lipid classification.

**Supplemental Data Set 4B**. Maximum normalized lipidomic data from wild-type, *pdf2*, *atml1*, and *gl2* mutants, as well as *EYFP:PDF2*, *EYFP:pdf2*$^{\Delta START}$, *EYFP:GL2*, and *EYFP:gl2*$^{L480P}$ transgenic lines in Pi sufficient (P+) and Pi limiting (P-) media. Related to **Figure 4** and **Extended Data Fig. 4**. Given are compound ID, lipid classification and normalized intensities for each sample. Averages were calculated from n = 4 or 5 biological replicates. Minimum and maximum values were normalized to 0.0 and 1.0, respectively, for visualization purposes in heat maps.

**Supplemental Data Set 4C**. Volcano plot calculations for lipidomic data from wild-type, *pdf2*, *atml1*, and *gl2* mutants, as well as *EYFP:PDF2*, *EYFP:pdf2*$^{\Delta START}$, *EYFP:GL2*, and *EYFP:gl2*$^{L480P}$ transgenic lines in Pi sufficient (P+) and Pi limiting (P-) media. Related to **Figure 4** and **Supplemental Figure 5**. Given are cluster name, m/z, retention time (RT), compound ID, lipid classification, adduct, and intensities for each sample. Averages and standard deviations were calculated from n = 4 or 5 biological replicates. FC values and corresponding log2 values are given for each of the mutants over WT. P-values were calculated from unpaired t-tests. T-test and FC were calculated only if data were available for three or more replicates. If a molecule was found in one or none of the samples in WT but in at 3 of 4 or 4 of 5 replicates in the mutant, the FC and P-value were assigned as 2 and 0.05, respectively. Color scheme: FC>2, light red; FC>4, red; FC<2, light blue; FC<4, blue (P-value <0.05).

**Supplementary Data Set 5A**. Lipidomics Mass Spectrometry Details: All Detected Peaks, Putative Metabolite Name Identified: Tandem Affinity Purification Experiment. Related to **Supplemental Data Set 1** and **Figure 1**.

**Supplemental Data Set 5B**. Lipidomics Mass Spectrometry Details: All Detected Peaks, Putative Metabolite Name Identified: Mutants Experiment. Related to **Supplemental Data Set 3**, **Supplemental Figure 2**, and **Supplemental Figure 3**.

**Supplemental Data Set 5C**. Lipidomics Mass Spectrometry Details: All Detected Peaks, Putative Metabolite Name Identified: Pi Limitation Experiment. Related to **Supplemental Data Set 4**, **Figure 4**, **Supplemental Figure 4** and **Supplemental Figure 5**.

### SUPPLEMENTAL REFERENCES


**Baumann, E., Lewald, J., Saedler, H., Schulz, B., and Wisman, E.** (1998). Successful PCR-based reverse genetic screens using an En-1-mutagenised Arabidopsis thaliana population generated via single-seed descent. Theor Appl Genet **97,** 729-734.

**Ishida, T., Hattori, S., Sano, R., Inoue, K., Shirano, Y., Hayashi, H., Shibata, D., Sato, S., Kato, T., Tabata, S., Okada, K., and Wada, T.** (2007). Arabidopsis TRANSPARENT TESTA GLABRA2 is directly regulated by R2R3 MYB transcription factors and is involved in regulation of GLABRA2 transcription in epidermal differentiation. Plant Cell **19,** 2531-2543.

**Kamata, N., Okada, H., Komeda, Y., and Takahashi, T.** (2013). Mutations in epidermis-specific HD-ZIP IV genes affect floral organ identity in Arabidopsis thaliana. Plant J.

**Khosla, A., Paper, J.M., Boehler, A.P., Bradley, A.M., Neumann, T.R., and Schrick, K.** (2014). HD-Zip Proteins GL2 and HDG11 Have Redundant Functions in Arabidopsis Trichomes, and GL2 Activates a Positive Feedback Loop via MYB23. Plant Cell **26,** 2184-2200.

**Li, M., Welti, R., and Wang, X.** (2006). Quantitative profiling of Arabidopsis polar glycerolipids in response to phosphorus starvation. Roles of phospholipases D zeta1 and D zeta2 in phosphatidylcholine hydrolysis and digalactosyldiacylglycerol accumulation in phosphorus-starved plants. Plant Physiol **142,** 750-761.

**Meyer, H.M., Teles, J., Formosa-Jordan, P., Refahi, Y., San-Bento, R., Ingram, G., Jonsson, H., Locke, J.C., and Roeder, A.H.** (2017). Fluctuations of the transcription factor ATML1 generate the pattern of giant cells in the Arabidopsis sepal. Elife **6**.

**Ogawa, E., Yamada, Y., Sezaki, N., Kosaka, S., Kondo, H., Kamata, N., Abe, M., Komeda, Y., and Takahashi, T.** (2015). ATML1 and PDF2 Play a Redundant and Essential Role in Arabidopsis Embryo Development. Plant Cell Physiol **56,** 1183-1192.

**Su, Y., Li, M., Guo, L., and Wang, X.** (2018). Different effects of phospholipase Dzeta2 and non-specific phospholipase C4 on lipid remodeling and root hair growth in Arabidopsis response to phosphate deficiency. Plant J **94,** 315-326.

**Wang, S., Kwak, S.H., Zeng, Q., Ellis, B.E., Chen, X.Y., Schiefelbein, J., and Chen, J.G.** (2007). TRICHOMELESS1 regulates trichome patterning by suppressing GLABRA1 in Arabidopsis. Development **134,** 3873-3882.

**Xia, J., Psychogios, N., Young, N., and Wishart, D.S.** (2009). MetaboAnalyst: a web server for metabolomic data analysis and interpretation. Nucleic Acids Res **37,** W652-660.
